# Suberin plasticity to developmental and exogenous cues is regulated by a set of MYB transcription factors

**DOI:** 10.1101/2021.01.27.428267

**Authors:** Vinay Shukla, Jian-Pu Han, Fabienne Cléard, Linnka Lefebvre- Legendre, Kay Gully, Paulina Flis, Alice Berhin, Tonni Grube Andersen, David E Salt, Christiane Nawrath, Marie Barberon

## Abstract

Suberin is a hydrophobic biopolymer that can be deposited at the periphery of cells, forming protective barriers against biotic and abiotic stress. In roots, suberin forms lamellae at the periphery of endodermal cells where it plays crucial roles in the control of water and mineral transport. Suberin formation is highly regulated by developmental and environmental cues. However, the mechanisms controlling its spatiotemporal regulation are poorly understood. Here, we show that endodermal suberin is regulated independently by developmental and exogenous signals to fine tune suberin deposition in roots. We found a set of four MYB transcription factors (MYB41, MYB53, MYB92 and MYB93), that are regulated by these two signals, and are sufficient to promote endodermal suberin. Mutation of these four transcription factors simultaneously through genome editing, lead to a dramatic reduction of suberin formation in response to both developmental and environmental signals. Most suberin mutants analyzed at physiological levels are also affected in another endodermal barrier made of lignin (Casparian strips), through a compensatory mechanism. Through the functional analysis of these four MYBs we generated plants allowing unbiased investigations of endodermal suberin function without accounting for confounding effects due to Casparian strip defects, and could unravel specific roles of suberin in nutrient homeostasis.

## Introduction

Plant roots form an inverted epithelium responsible for the selective acquisition of water and nutrients present in the soil. When entering the root, water and nutrients need to be radially transported from the root periphery to the central vasculature in order to be loaded to the xylem vessels and distributed to the plant organs. This can be achieved through three different transport scenarios: symplastic, apoplastic or transcellular (1, 2). The endodermis, the innermost cortical cell layer surrounding the central vasculature, plays a particularly important role in these transport routes as it forms barriers for the free diffusion of water and nutrients. These barriers are formed in two sequential differentiation stages with first, the formation of Casparian strips (CS), ring-like structures made of lignin forming an apoplastic barrier (3-5), and then suberin lamellae deposited as secondary cell walls around endodermal cells forming a diffusion barrier for the transcellular pathway (5-7). Recent efforts studying mutants and lines affected for CS and/or endodermal suberin in *Arabidopsis thaliana* and in rice allowed to demonstrate that both barriers play crucial roles in nutrient acquisition and homeostasis (6, 8-13). Yet, the role of suberin in nutrient transport is still poorly understood and, in the absence of mutants with constitutive strong reduction in suberization, is mainly corroborated by the analysis of a synthetic suberin-deficient line (artificially expressing in the endodermis the cutinase CDEF1, CUTICLE DESTRUCTING FACTOR1, to degrade suberin) (6, 11, 14). To complicate matters, most known enhanced suberin mutants are actually Casparian strip defective mutants, with ectopic endodermal lignification and suberization acting as compensation (9-11). This syndrome occurs in response to Casparian strip defects and is triggered through the endodermal integrity control system consisting of the Leucine-rich-repeat Receptor-like Kinase, SGN3/GSO1 (SCHENGEN3/GASSHO1) and its ligands CIF1/2 (CASPARIAN STRIP INTEGRITY FACTORS 1/2) (12, 15-19). Suberin, however, is not only regulated by endogenous developmental factors surveilling Casparian strip integrity. Pointing towards a very central role of suberin in plant adaptation to their environment, endodermal suberization is also highly regulated by nutrient availability and the hormones ethylene and ABA (abscisic acid) (6, 14, 20-25), as well as during biotic interactions (25-28). How suberin is regulated in response to developmental and exogenous clues remains poorly understood. Recently, several transcription factors were shown to be sufficient to induce ectopic suberin formation when ectopically overexpressed and for some to directly activate the expression of suberin biosynthesis genes (29-33). Suggesting a potential role in controlling endodermal suberization the transcription factors *MYB39, MYB93* (*MYeloBlastosis* family of transcription factors) and *ANAC046* (*Arabidopsis thaliana NAM/ATAF/CUC protein*) were shown to be constitutively expressed in the endodermis and *MYB41* to be expressed in the endodermis in response to ABA or salt, two conditions known to induce suberization (29, 31, 32, 34). However, in the absence of clear suberin phenotypes associated with loss of function, their actual role in endodermal suberin formation and its regulation remains unclear.

Here by combining epistasis and pharmacological experiments, we demonstrated that suberin is regulated independently by the SGN3/CIFs pathway and ABA (previously shown to control suberin induction in response to nutritional stresses). We next undertook a systematic gene expression analysis and identified four endodermal MYB transcription factors (MYB41, MYB53, MYB92 and MYB93) acting downstream of SGN3/CIFs and ABA signaling, in the endodermis. These transcription factors are sufficient to induce suberin biosynthesis in the endodermis. Moreover, we generated a quadruple mutant by CRISPR/Cas9 and could show that these transcription factors are necessary to form endodermal suberin and to induce suberization in response to developmental and exogenous signaling. Our work developed plants specifically and strongly impaired in endodermal suberin allowing us not only to probe the regulatory mechanisms of suberin formation but also to characterize the specific function of suberin in nutrient homeostasis.

## Results

### Suberin is induced by ABA and SGN3/CIFs independently

Suberin formation can be induced in response to nutrient availability through ABA and in response to Casparian strip defects through the receptor SGN3/GSO1 and its ligands CIF1/2 (6, 12, 16, 17). In order to investigate the underlying molecular mechanism controlling ectopic suberization it was important to establish if ABA and SGN3/CIFs have a similar effect on suberin formation. We compared the effects of exogenous applications of the hormone ABA and the peptide CIF2 on root suberization. To this end we used the suberin biosynthesis reporter line *GPAT5::mCitrine-SYP122* (driving the expression of a fluorescently tagged plasma membrane anchor protein under the control of the promoter of the suberin biosynthesis gene *Glycerol-3-Phosphate Acyl Transferase5)* and whole-mount suberin staining using Fluorol Yellow (FY) (Fig. 1A-C). In untreated roots, we observed a typical pattern of suberin formation (5, 6, 35) with a non-suberized zone (state I of endodermal differentiation with Casparian strips) followed by a suberizing zone where only patches of endodermal cells are suberized (patchy zone) and finally a fully suberized zone (Fig. 1A-C). Exogenous treatments with ABA or CIF2 peptide led to ectopic suberin formation at the proximity of the root tip, without a patchy zone between non-suberized and fully suberized zones (Fig. 1A-C) with ABA additionally inducing further suberization in the fully differentiated endodermis and in cortical cells, as described before (6). Both treatments had the same effect on the onset of endodermal suberization, which begs the question whether these two signals are converging on the same mechanism. This has been addressed before and independent works on this suggest either an interaction between ABA and developmental signals (36), or, on the contrary, an independence (14). In light of these contradictions, we decided to clarify the relation between ABA and SGN/CIFs as signals controlling endodermal suberization. We first tested if the CIF1/2 receptor SGN3/GSO1 was needed for ABA-dependent suberization. We used the CIF-insensitive mutant *sgn3* and observed no difference between ABA induced ectopic endodermal suberization in WT plants and in *sgn3* mutants (Fig. 1C,D). This hints to ABA signaling being active either downstream or independent of the SGN3/CIFs pathway. To elucidate this further, we assessed if exogenous CIF2 application can induce suberization in absence of active ABA signaling. We used the previously described *ELTP::abi1-1* line, where ABA signaling is inhibited in the endodermis by expressing the dominant-negative *abi1-1* (*aba insensitive 1*) allele specifically in the endodermis using the *ELTP/EDA4* promoter (*Endodermal Lipid Transfer Protein* / *EMBRYO SAC DEVELOPMENT ARREST 4*) (6). As previously reported for *ELTP::abi1-1* plants, endodermal suberization was severely delayed in non-stressed conditions (6, 35), but CIF2 application was able to induce suberin formation similarly to the response observed in WT plants (Fig. 1E). This indicates that ABA signaling is not acting downstream of SGN3/CIFs and that both pathways control suberization independently. To strengthen this conclusion, we tested the role of ABA signaling in the SGN3/CIFs-dependent enhanced suberin phenotype observed in the Casparian strip defective mutant *esb1* (*enhanced suberin 1*) (9, 12). We first expressed *ELTP::abi1-1* in *esb1* mutant background and observed an enhanced suberin phenotype independent of endodermal ABA signaling (Fig. S1A), confirming previous analysis (14). Next, we confirmed this observation by pharmacological interference with ABA biosynthesis using fluridone (an herbicide blocking the carotenoid biosynthesis indirectly and thus lowering the amount of ABA), widely used as an ABA biosynthesis inhibitor (37-39). In presence of fluridone, suberin is highly reduced in WT plants but the *esb1* mutant still displays its enhanced suberin phenotype (Fig. S1B). Altogether these data demonstrate that ABA and SGN3/CIFs pathways can induce ectopic endodermal suberization independently.

**Figure 1.**
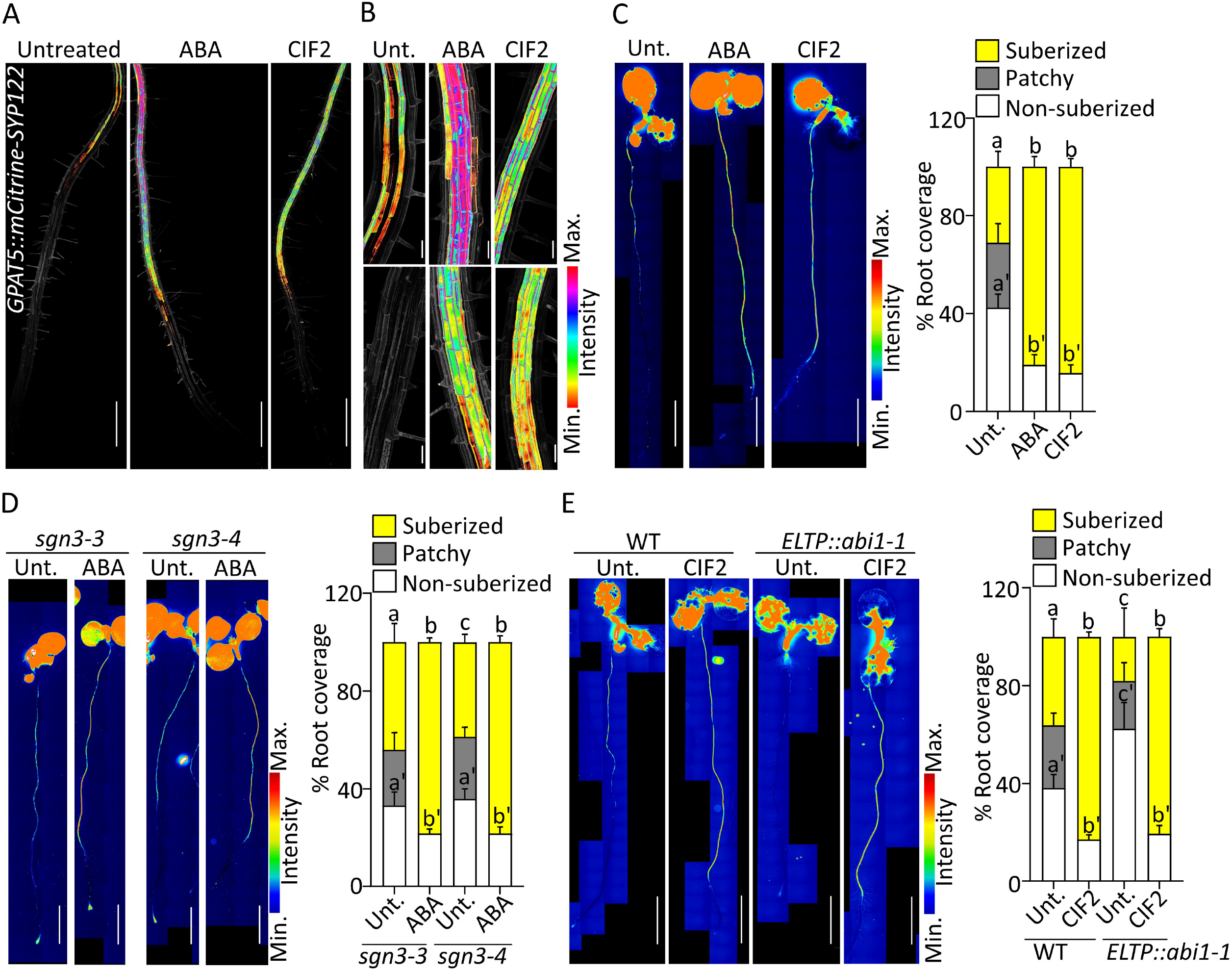
Independent suberin induction by ABA and SGN3/CIF. *(A,B) GPAT5::mCITRINE-SYP 122* expression (presented as LUT) in untreated (Unt.) or plants treated with 1 µM ABA or 1 µM CIF2 for 16 h. Propidium iodide (PI, in grey) was used to highlight cells. Pictures are presented as maximum intensity Z projections. *(A)* Observations from root tip to 4-5 mm. Scale bars, 500 µm. *(B)* Zoomed pictures from *A* in the patchy *(Upper)* and non-suberized zones *(Lower)* respectively. Scale bars, 50 µm. *(C-E)* Fluorol Yellow (FY) staining for suberin. Whole-mount staining in full seedlings *(Left panels)* and quantifications of suberin pattern along the root *(Right panels)*, n 10, error bars, standard deviation, different letters indicate significant differences between conditions *(P* < 0.05). Scale bars, 2 mm. (C) FY staining for suberin in Unt. or treated WT plants with 1 µM ABA or 1 µM CIF2 for 16 h. *(D)* FY staining in *sgn3-3* and *sgn3-4* mutants Unt. or treated with 1 M ABA for 16 h. *(E)* FY stainin for WT and *ELTP::abil-1* Unt. or treated with 1 µM CIF2 for 16 h.

### MYB41 is a primary response factor to suberin inducing signals

Next, we aimed to identify transcription factors controlling endodermal suberization downstream of ABA and SGN3/CIFs. Several MYB transcription factors - *MYB9, MYB39, MYB41, MYB53, MYB92, MYB93* and *MYB107 -* have been shown in transient assays in *Nicotiana benthamiana* leaves, ectopic overexpression in whole plant and/or yeast one-hybrid experiments to be able to activate suberin biosynthesis (29-33). Among them, MYB9 and MYB107 were shown to control suberin deposition in seed coats (40, 41). Very recently, MYB39 has been proposed as a regulator of endodermal suberization (31). However, the *myb39* mutant showed only a partial delay in endodermal suberization suggesting the involvement of other transcriptional regulators. Moreover, the primary factors regulating suberin biosynthesis in response to exogenous and developmental cues were still unknown. To identify such factors, we narrowed our search to the *MYBs* whose expression was induced by both ABA and the SGN3/CIFs pathway. We mined publicly available transcriptomes in seedlings treated with ABA for 1h and 3h (42) and roots treated with CIF2 peptide for 2h and 8h (17) and found a moderate response of all the selected *MYBs* (*i*.*e. MYB39, 53, 92* and *93*), with the exception of *MYB41* whose expression responded the fastest and strongest to either stimuli (Fig. S2A). To validate this observation, we performed a time-course experiment for transcript profiling of roots after 3h and 6h of ABA or CIF2 applications in our growth conditions (Fig. 2A). We confirmed that *MYB41* was indeed the primary responsive factor for either stimuli. The other factors *MYB53, MYB92* and *MYB93* were also induced by both stimuli but at a lower level while *MYB39* expression was reduced after ABA and CIF2 applications in our conditions and was therefore not investigated further in this study (Fig. 2A).

**Figure 2.**
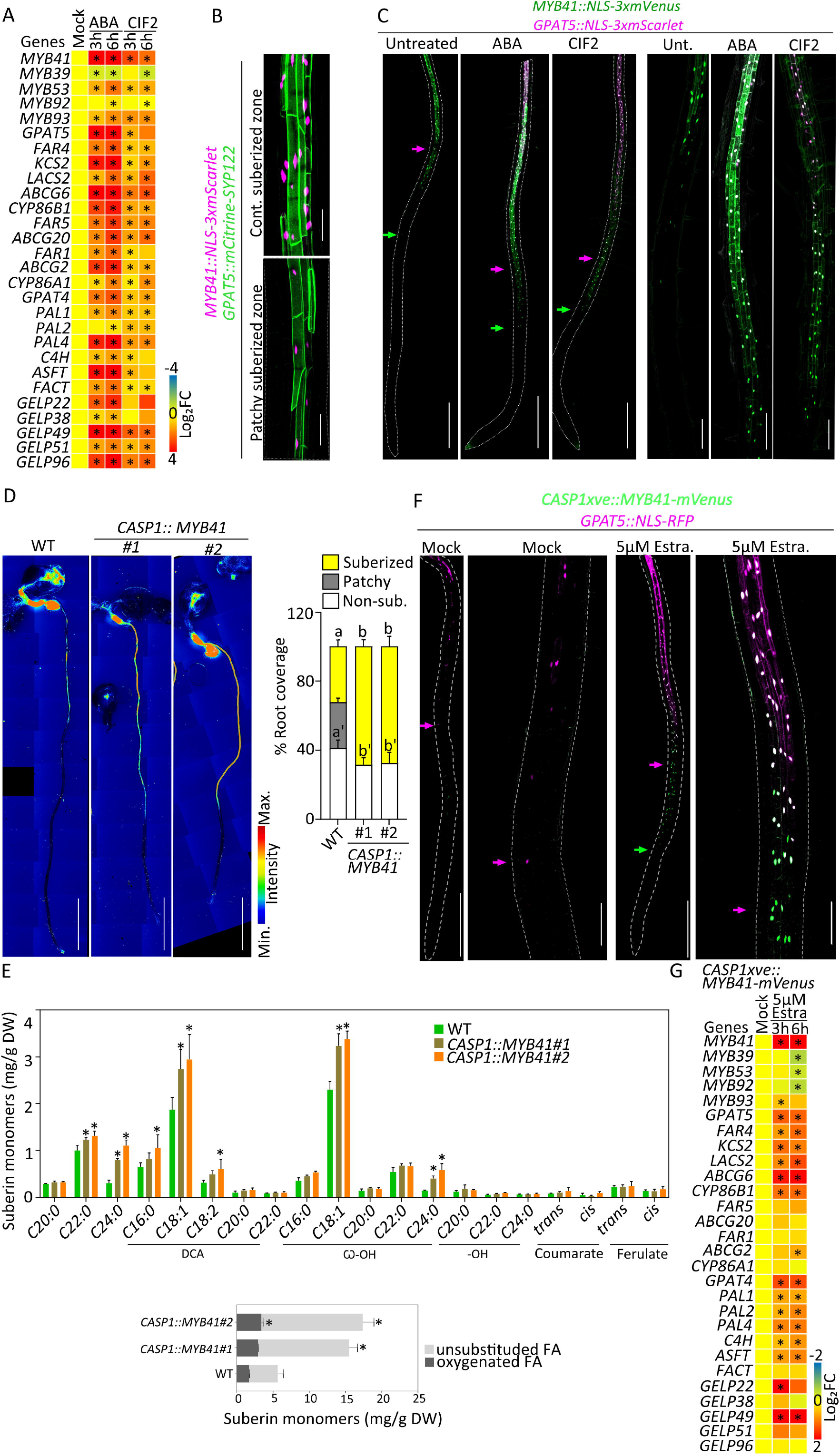
MYB41, an endodermal transcription factor, inducing suberin biosynthesis. *(A)* Relative expression levels of the candidate *MYBs* and suberin biosynthesis and polymerization genes in WT roots treated with 1 µM ABA or 1 µM CIF2 for 3 and 6 h (n = 4 pools of25-30 roots). Results are presented as fold changes compared to the mock condition. Numeric values are presented in Table S3. Asterisks indicate statistically significant differences *(P* < 0.05). *(B)* Dual reporter for *GPAT5::mCitrine-SYP122* and *MYB41::NLS-3xmScarlet* under untreated condition. Pictures are presented as maximum intensity Z projections taken in the zone of continuous suberization *(Upper)* and patchy suberization *(Lower)*. Scale bars, 50 µm. (C) Dual reporter for *GPAT5::NLS-3xmScarlet* and *MYB41::NLS-3xmVenus* upon 16 h treatments with 1 µM ABA or 1 µM CIF2. Pictures are presented as maximum intensity Z projections from the root tip to 4-5 mm *(Left panels)* and zoomed views *(Righ panels)* corresponding to the zone prior to suberization in untreated conditions. Arrows highlight the onset of *MYB41* (green) and *GPAT5* (magenta) expression. Scale bars, 500 µm. *(D)* FY staining of WT and two independent *CASPJ::MYB41* lines. Whole-mount staining *(Left panels)* and quantifications of suberin pattern are presented *(Right panel)*, n ≥ 10, error bars, standard deviation, different letters indicate significant differences between conditions *(P* < 0.05). Scale bars, 2 mm. *(E)* Polyester composition in 5-day-old roots of WT and two independent *CASP1::MYB41* lines. Individual suberin monomer content (*Upper panel)* and the corresponding total amount of unsubstituted and oxygenated fatty acids *(Lower panel)* are presented. Data correspond to mean; error bars, standard deviation (n = 4 pools of 200-300 roots). FA, fatty acid; DCA, dicarboxylic fatty acid; co - OH, co-hydroxy fatty acid; DW, dry weight. Asterisks represent statistical significant differences compared to WT *(P* < 0.05). Results from WT are also shown in Fig. 4B. *(F) CASPlxve::MYB41-mVenus* in *GPAT5::NLS-RFP* background for mock and 5 µM Estradiol (Estra.) treatment for 16 h . Pictures are presented as Z projections from the root tip to 4-5 mm *(Left)* and zoomed views *(Right)* taken in the corresponding zone for patchy suberization in mock conditions. Arrows highlight the onset of *MYB41* (green) and *GPAT5* (magenta) expressions. Scale bars, 500 µm *(Left)* and 100 µm *(Right)*. (G) Relative expression levels of the *MYBs* candidates and suberin biosynthesis and polymerization genes in the roots of *CASPlxve::MYB41-mVenus* treated with 5 µM Estradiol for 3 and 6 h (n = 4 pools of 25-30 roots). Results are presented as fold changes compared to the mock condition. Numeric values are presented in Table S3. Asterisks indicate statistically significant differences *(P* < 0.05). CIF2 for 16 h. Different letters indicate significant differences between conditions for a given genotype *(P* < 0.05).

Since *MYB41* reacted most prominently of all MYBs from transcript profiling, we focused on this factor as the primary candidate for controlling endodermal suberization and its induction by ABA and SGN3/CIFs. To test the spatiotemporal response of *MYB41* upon ABA and CIF2 applications, we developed a transcriptional reporter line, *MYB41::NLS-3xmVenus* for live imaging in roots. We found that *MYB41* was specifically expressed in the differentiated endodermis, matching the tissue-specificity of the suberin biosynthesis reporter *GPAT5* (Fig. 2B). Applications of ABA and CIF2 further validated the transcriptional response observed in previous experiments of mRNA profiling as we observed a strong activation of *MYB41* promoter activity in the endodermis with an ectopic expression close to the root tip (Fig. S2B, C). We combined this reporter with *GPAT5::NLS-3xmScarlet-I* to generate a dual-reporter for *MYB41* and *GPAT5* promoter activity and observed in untreated conditions, that *MYB41* expression preceded the expression of *GPAT5* in the endodermis, positioning *MYB41* in spatio-temporal context for regulating endodermal suberization (Fig. 2C). Upon ABA and CIF2 applications, *MYB41* expression was strongly induced, and its expression pattern extended to the proximity of the root tip. Since *MYB41* expression always preceded *GPAT5* spatiotemporal expression (Fig. 2C, Fig. S2D-F), this could be indicative of *MYB41* controlling the suberin biosynthesis machinery in the endodermis.

We then tested if MYB41 activity was sufficient to induce endodermal suberization. To this end, we used the endodermis-specific promoter *CASP1* (expressed in the differentiating endodermis before suberization) to drive *MYB41* expression. FY staining of *CASP1::MYB41* transgenic lines showed ectopic endodermal suberization closer to the root tip, demonstrating that *MYB41* expression was sufficient for induction of endodermal suberization (Fig. 2D). We confirmed this observation by performing chemical analysis of suberin content in the roots of WT and two independent *CASP1::MYB41* lines. We found that both *CASP1::MYB41* lines displayed excess of suberin monomers with an increase of ∼140% for line #3 and ∼170% for line #7 compared to WT roots (Fig. 2E). We simultaneously tested the same in rather synthetic manner by expressing a functional *MYB41-mVenus* under a chemically inducible endodermal promoter, *CASP1xve* in WT (Fig. S2G) and *GPAT5::NLS-RFP* reporter backgrounds. Importantly after estradiol induction, we could observe a transient accumulation of MYB41-mVenus in endodermal cells followed by the induction of *GPAT5* promoter activity, corroborating our supposition that MYB41 can induced suberin biosynthesis in the endodermis (Fig. 2F, Fig. S2H, J). In order to verify if this conditional induction of MYB41 was able to induce the rest of the suberin biosynthetic pathway, we measured the transcript levels of suberin biosynthesis genes and the other *MYBs* of interest. A short treatment of 3h with estradiol was enough to strongly induce *MYB41* expression as well as nearly all the genes involved in suberin biosynthesis, including the recently characterized *GELPs* (*GDSL-type Esterase/Lipases*) (43) coding for enzymes involved in the polymerization of suberin monomers in the cell wall (Fig. 2G). Surprisingly we could also observe an increased expression for *ASFT, PAL1, PAL2, PAL4* and *C4H* while no significant increase in ferulate content was detected (Fig. 2E,G). In addition, we observed that MYB41 did not induce the expression of most other *MYBs* studied with only a transient induction of *MYB93* expression and a reduction of the expression of *MYB39, MYB53* and *MYB92* after 6h estradiol induction (Fig. 2G). This reduction of *MYB39, MYB53* and *MYB92* expression being observed only after 6h while the expression of genes involved in suberin biosynthesis was already induced after 3h likely reflects a compensatory effect.

### A set of four MYBs control suberin biosynthesis and regulation

After establishing that MYB41 was sufficient to induce endodermal suberization, we wondered whether it was also required to establish the endodermal suberin pattern observed under unstressed conditions in wildtype plants. We generated two CRISPR alleles of *MYB41*. The CRISPR mutants, *myb41_c1* was obtained by a nearly full deletion of the *MYB41* coding region and *myb41_c2* was obtained by introducing a one-base-pair frame shift in the beginning of the third and longest exon of *MYB41* gene (Fig. S3A). To confirm the protein inactivity from the point mutation generated in *myb41_c2*, the mutated *MYB41_c2* cDNA was cloned and expressed in plants using the *CASP1* promoter and, unlike the unmutated MYB41 cDNA, was unable to induce suberization (Fig. S3B). Unexpectedly, FY staining showed that suberin deposition in the endodermis was unaffected in these two CRISPR mutants (Fig. 3A). Moreover, ABA or CIF2 treatment induced ectopic suberization in *myb41_c1 and myb41_c2* mutants virtually indistinguishable from WT plants (Fig. 3A). These observations led us to consider that though *MYB41* was the primary responsive factor to ABA and CIF2 and is sufficient to induce suberization, other functionally redundant MYBs are probably compensating in its absence.

**Figure 3.**
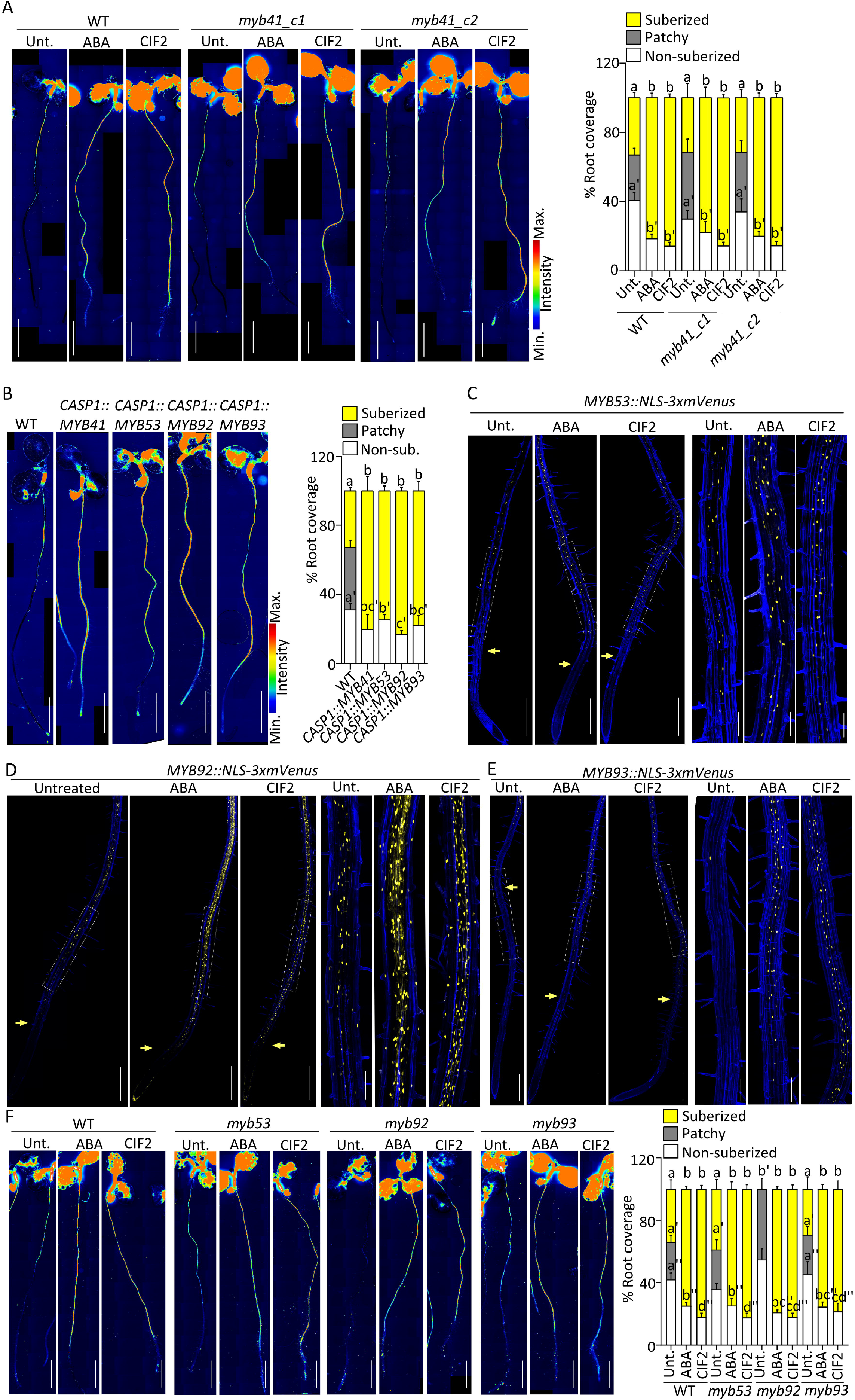
*MYB53, MYB92* and *MYB93* regulation and function in suberin induction. *(A,B, F)* Fluorol Yellow (FY) staining for suberin. Whole-mount staining *(Left panels)* and quantifications of suberin pattern along the root *(Right panels)*, n ≥ 10, error bars, standard deviation. Scale bars, 2 mm. *(A)* FY staining of WT and two *myb41* CRISPR mutant alleles, *myb4l_cl* and *myb4l_c2* untreated or treated with 1 µM ABA or 1 µM CIF2 for 16 h. Different letters indicate significant differences between conditions for a given genotype *(P* < 0.05). *(B)* FY staining of WT, *CASPl::MYB41, CASPJ::MYB53, CASPJ::MYB92* and *CASPJ::MYB93* lines. Different letters indicate significant differences between conditions *(P* < 0.05). (C) *MYB53::NLS-3xmVenus, (D) MYB92::NLS-3xmVenus* and *(E) MYB93::NLS-3xmVenus* expression (in yellow) untreated or after treatments with 1 µM ABA or 1 µM CIF2 for 16 h. (*C-E)* Pictures are presented as maximum intensity Z projections taken from the root tip to 4-5 mm *(Left panels)* with zoomed views corresponding to the zone of patchy suberization in untreated plants *(Right panels)*. Arrows highlight the onset of expression. Propidium iodide (PI, in blue) was used to highlight cells. Scale bars, 500 µm *(Left)* and 125 µm *(Right)*. Quantifications are shown in Fig. S3D-F. (F) FY staining of WT, *myb53, myb92* and *myb93* mutant untreated or treated with 1 µM ABA or 1 µM CIF2 for 16 h. Different letters indicate significant differences between conditions for a given genotype *(P* < 0.05)

To identify other MYBs that might be active in the *myb41* mutants, we compared the transcript levels of the other *MYB* candidates in WT and *myb41_c1* mutant backgrounds and found that the mRNA levels of *MYB39, MYB53, MYB92*, and *MYB93* were slightly higher in *myb41_c1* compared to the WT in untreated conditions (Fig. S3C). Upon ABA and CIF applications, all the *MYBs* except *MYB39*, were further induced in both backgrounds. Therefore, we decided to additionally characterize *MYB53, MYB92* and *MYB93* in endodermal suberization. Similarly to MYB41, the selected MYB candidates, MYB53, MYB92 and MYB93, are able to induce ectopic endodermal suberization. FY staining of the lines *CASP1::MYB53, CASP1::MYB92* and *CASP1::MYB93* along with *CASP1::MYB41*, clearly showed that endodermis specific precocious expression of any of these *MYB*s was sufficient to induce suberization close to the root tip similarly to *MYB41* (Fig. 3B). Next, we wondered which one of these *MYBs* were expressed in the endodermis and generated promoter-reporter lines for *MYB53, MYB92* and *MYB93* driving the expression of *NLS-3xmVenus*. We found that, similarly to *MYB41, MYB53, MYB92* and *MYB93* were also expressed in unstressed conditions in the endodermis with *MYB53* and *MYB92* expressed from the differentiated zone (Fig. 3C-E). However, *MYB93* expression was observed only in few cells, most likely in the endodermis above the lateral root primordia (Fig. 3E) as it was previously described (34). Additionally, *MYB53* and *MYB92* were also expressed in few isolated cortical and epidermal cells in unstressed conditions (Fig. 3C-D). Importantly, all these promoters promptly responded to ABA and CIF2 resulting in a higher expression, with the expression of *MYB93* extending to the endodermal cells close to the root tip as observed for *MYB41* (Fig. 3C-D, Fig. S3D-F). Taken together, we concluded that not a single MYB, but rather a group of MYB transcription factors are likely controlling suberin biosynthesis and regulation in the endodermis. To test this hypothesis we characterized with FY staining the pattern of suberin deposition in *myb53, myb92* and *myb93* mutants in unstressed condition and in response to ABA and CIF2 treatments (Fig. 3F). While *myb53* and *myb93* displayed no suberin phenotype in all conditions tested, *myb92* showed a significant delay in suberin deposition in unstressed condition with suberin deposited later from the root tip and only as patches of suberized cells (Fig. 3F). However, even in *myb92* mutant ABA or CIF2 treatment induced ectopic suberization similarly to the effect observed in WT plants (Fig. 3F). We therefore decided to mutate all these 4 *MYBs* simultaneously by using multiplexed CRISPR technology (44) introducing frame shifts leading to loss of function in *MYB53, MYB92* and *MYB93* in the mutant background, *myb41_c2* (Fig. S4A-B). After FY staining of the resulting quadruple mutant – *myb41-myb53-myb92-myb93* (*quad-myb*) we observed nearly a total absence of suberin in roots (Fig. 4A). Importantly, suberin induction by ABA and CIF2 treatments was also severely compromised in the *quad-myb* mutant where almost no induction was observed after 3 or 6 h and only a weak effect was observed after 16h (Fig. 4A, Fig. S4C). We confirmed this observation by performing chemical analysis of the suberin content in the roots of the *quad-myb* mutant compared to WT roots and found a strong reduction of both aliphatic and aromatic monomers (Fig. 4B). Dicarboxylic acids and ω-hydroxy acids were nearly absent with ∼90% reductions, while reductions in fatty alcohols ranged from 20 to 80%. Ferulate was reduced by ∼80% and coumarate showed ∼50% reduction compared to WT. On average, the quadruple mutant showed an overall ∼78% decrease in suberin monomers compared to WT (Fig. 4B). We also quantified the mRNA levels of suberin biosynthesis and polymerization genes and found that most were accumulating at lower levels, especially the genes involved in fatty acid pathway and polymerization (Fig. 4C). The expression of genes involved in the phenylpropanoid biosynthesis were also affected with *PAL2* and *PAL4* expressed at higher levels but without leading to an increase of aromatic suberin components (Fig. 4B-C). These genes being also expressed at higher levels in *CASP1xve::MYB41* after estradiol induction, their change in expression (Fig. 2G) is not directly affected by changes in *MYBs* expression. Altogether, this strongly supports a central role of the 4 transcription factors MYB41, MYB53, MYB92 and MYB93 in the control of suberin biosynthesis but also for its regulation by the two main signals inducing endodermal suberization.

**Figure 4.**
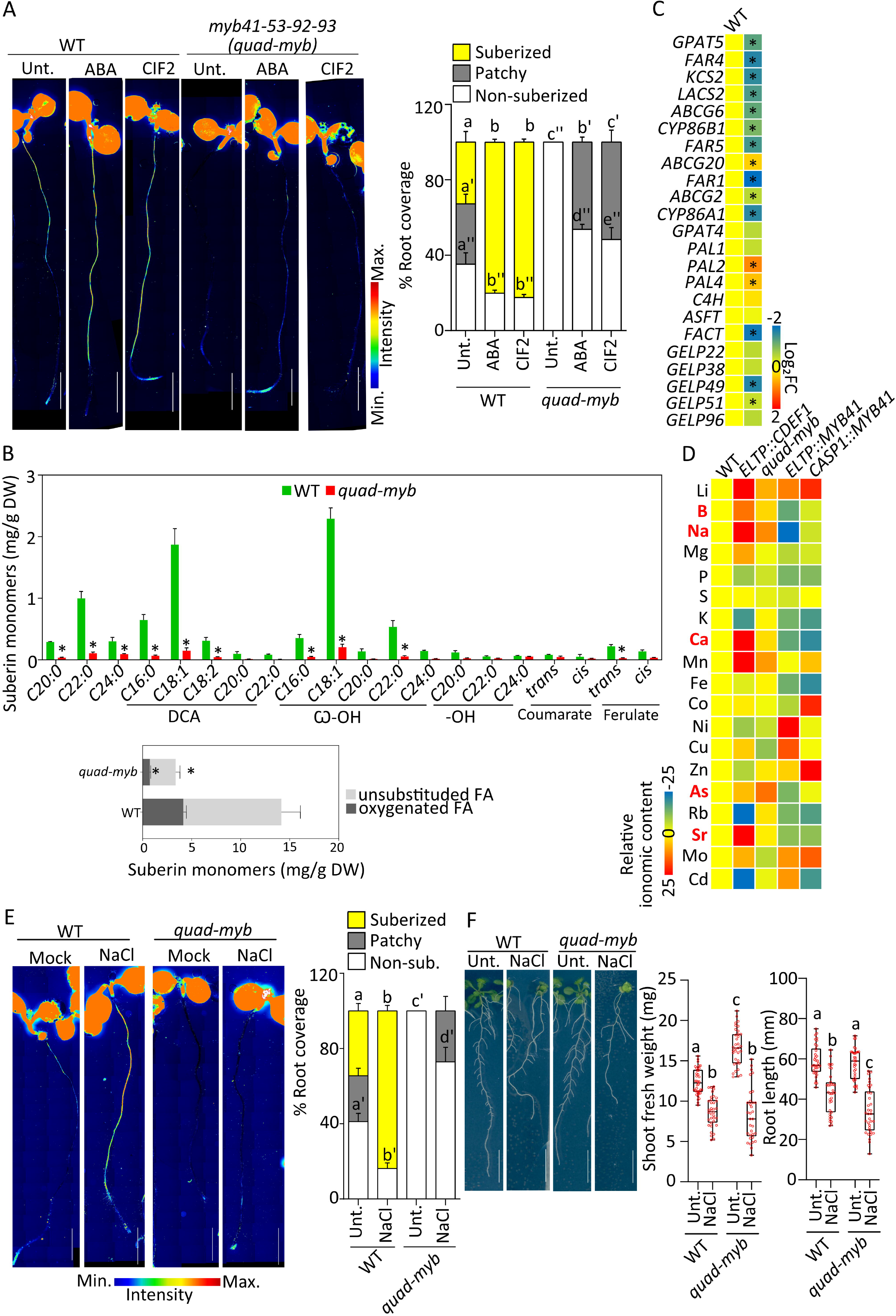
Impaired suberin formation and regulation in *quad-myb* mutant associate with nutritional changes. *(A,E)* Fluorol Yellow (FY) staining for suberin. Whole-mount staining *(Left panels)* and quantifications of suberin pattern along the root *(Right panels)*, n ≥ 10, error bars, standard deviation, different letters indicate significant differences between conditions (P < 0.05). Scale bars, 2 mm. (A) FY staining ofWT and *myb41-myb53-myb92-myb93 (quad-myb)* mutant untreated or treated with 1 µM ABA or 1 µM CIF2 for 16 h. (B) Polyester composition in 5-day old roots ofWT and two independent *CASP1::MYB4l* lines. Individual suberin monomer content (*Upper panel)* and the corresponding total amount of unsubstituted and oxygenated fatty acids *(Lower panel)* are presented. Data are represented as mean; error bars, standard deviation (n = 4 pools of 200-300 roots). FA, fatty acid; DCA, dicarboxylic fatty acid; ro -OH, ro-hydroxy fatty acid; DW, dry weight. Asterisks represent statistically significant differences (P < 0.05). Results from WT are also shown in Fig. 2E. (C) Relative expression levels of the suberin biosynthesis and polymerization genes in the roots of *quad-myb* mutant compared to WT (n = 4 pools of 25-30 roots). Results are presented as fold changes compared to theWT. Numeric values are presented in Table S3. Asterisks indicate statistical significance (P < 0.05). (D) Ionomic profiling of leaves of WT, *ELTP::CDEFJ, quad-myb, ELTP::MYB41* and *CASPl::MYB41* plants. Elements were determined by ICP-MS. Results are presented as average fold changes compared to theWT. Results from individual experiments are presented in Fig. S5D and numeric values in Table S4. (E) FY staining ofWT and *quad-myb* mutant untreated or treated with 75 mM NaCl for 16 h. (F) NaCl effect on *quad myb* root shoot weight and root length. WT and *quad-myb* plants were grown 3 days after which they were treated different letters indicate significant differences between conditions *(P* < 0.05). Scale bars, 10 mm. with 75 mM NaCl for next 10 days. Results are presented as pictures of plants at the end of day 13 *(Left panels)* and as quantifications *(Right panels)*. Data are presented as box plots with individual values overlaid, n ≥ 30, error bars, different letters indicate significant differences between conditions (*p* < 0.05). Scale bars, 10mm.

### MYB-dependent suberization reveals specific roles of suberin in nutrient homeostasis

Having identified a set of four MYB transcription factors playing a central role in controlling suberin biosynthesis and regulation we set out to use these MYBs as tools to manipulate endodermal suberin specifically. Indeed, although suberin is known to play important roles for nutrient homeostasis we often cannot distinguish its role from Casparian strips. Previous efforts relied on mutants such as *esb1, casp1casp3*, or *myb36* with pleiotropic endodermal defects in Casparian strips formation (9, 10, 14). Here we generated *MYB41* overexpressing plants as potentially interesting tools to specifically enhance endodermal suberin in plants. To our surprise, plants overexpressing *MYB41* under the *CASP1* promoter displayed interrupted Casparian strips accompanied by a delay in the establishment of their apoplastic barrier (Fig. S4D-E) similar to the defects observed in *esb1* or *casp1casp3* mutants (9, 12). This may indicate that a precocious suberin formation (concomitant with Casparian strip formation) interferes with Casparian strip formation. Using the *ELTP* promoter, whose endodermal expression is much weaker than *CASP1* and is not affected by ABA and CIF2 (Fig. S4F-G), to trigger *MYB41* expression, we observed ectopic suberin formation close to the root tip in the corresponding plants (Fig. S4H) yet, importantly, without affecting Casparian strips and the establishment of the apoplastic barrier (Fig. S4D-E). We therefore have now access to dominant genetic tools to either enhance suberin specifically (*ELTP::MYB41*), or together with Casparian strip defects (*CASP1::MYB41*). In parallel we have generated in this study a novel mutant, *quad-myb*, with a dramatic reduction of suberin and observed no Casparian strip defects and no delay in the establishment of the apoplastic barrier (Fig. S4D-E). We therefore studied and compared the suberin-only affected plants *ELTP::MYB41*, with the non-suberized *quad-myb* mutant as well as with the suberin and Casparian strip affected *CASP1::MYB41* in order to understand better the consequences of reduced or enhanced endodermal suberin, independent or coinciding with Casparian strip defects. All plants generated were carefully studied for their growth and development in plates and in soil conditions. The corresponding seedlings were indistinguishable at early stages of development, where most of histological, chemical, expression and ionomic analysis where performed (Fig. S5C). At later stages we observed minor changes in the primary root length, although the number and length of lateral roots was highly reduced in both *ELTP::MYB41* and *CASP1::MYB41* lines and increased in the *ELTP::CDEF1* line (Fig. S5A). The *quad-myb* mutant was slightly affected in primary root length but not for lateral roots in our conditions (Fig. S5A). These changes in root architecture could be associated with enhanced or reduced suberization affecting directly root development, as it was previously suggested (11), or indirectly as a consequence of changes in nutrient acquisition. In soil, after 2 to 3 weeks of growth we could observe that all genotypes were comparable (Fig. S5B). Therefore, specific manipulation of endodermal suberin had no dramatic consequences on plant growth and development allowing further physiological characterization of the corresponding plants.

To study the consequences of endodermal suberin manipulation for nutrient homeostasis, we performed inductively coupled plasma-mass spectrometry (ICP-MS) for elemental profiling in leaves (Fig. 4D, Fig. S5D, Table S4). Confirming previous analysis in independent growth conditions (6), *ELTP::CDEF1* leaves accumulated arsenic, lithium, magnesium and sodium at higher levels and potassium and rubidium at lower levels compared to WT plants. We observed additional ionomic changes in our growth conditions with a higher accumulation of boron, calcium, manganese, strontium and molybdenum and a lower accumulation of phosphorous, iron, zinc and cadmium in *ELTP::CDEF1* compared to WT. The *quad-myb* mutant displayed also multiple ionomic changes compared to WT plants with similarities to several changes observed in *ELTP::CDEF1* although more moderately (Fig. 4D, Fig. S5D, Table S4). Among all these ionomic changes the levels of lithium, boron, sodium, calcium, manganese, arsenic and strontium were found higher and the levels of phosphorous, nickel and cadmium were found lower in both genotypes and could therefore be directly associated with an absence of endodermal suberin. On the other hand, *ELTP::MYB41* and *CASP1::MYB41* lines displayed more differences, with manganese, cobalt, nickel, copper, and cadmium accumulating in opposite manners (Fig. 4D). However, lithium and zinc accumulated at higher levels and boron, sodium, magnesium, phosphorous, potassium, calcium, iron, arsenic, rubidium, and strontium accumulated at lower levels in both *ELTP::MYB41* and *CASP1::MYB41* lines. To identify elements potentially directly affected by suberin among all these changes we selected the elements commonly affected in the suberin deficient *quad-myb* and *ELTP::CDEF1* plants and oppositely, but commonly, affected in the enhanced suberin *ELTP::MYB41* and *CASP1::MYB41* plants. Following this rational we found only few elements following this trend with boron, sodium, calcium, arsenic and strontium accumulating at higher levels in suberin deficient plants and at lower levels in enhanced suberin plants (Fig. 4D). Importantly these ionomic changes were not explained by compensations in the expression of genes encoding transporters (Fig. S5E). Among these elements, calcium and sodium were previously proposed to be directly affected by endodermal suberin and plants with reduced suberization were shown to accumulate these elements at higher levels (6, 11, 22, 45-47). In the context of sodium, endodermal suberin induction in response to salt stress was proposed to represents a protective mechanism against sodium entrance in plants. We set out to further test this hypothesis with the *quad-myb* mutant by studying its response to salt. First, we performed suberin staining on *quad-myb* treated with NaCl and observed that while this treatment induced suberization close to the root tip in WT plants like previously described (6), the *quad-myb* mutant was almost not responding (Fig. 4E). Next, we tested the tolerance of the *quad-myb* mutant to a mild salt treatment. When considering shoot weight and root length, *quad-myb* plants were significantly more reduced compared to WT plants when growing in presence of salt (Fig. 4F) to degrees similar to what was described before for *ELTP::CDEF1* and the quintuple *gelp22-38-49-51-96* mutant (6, 43). Combined, our results further support the central role of suberin in plant adaptation to the presence of salt.

## Discussion

Suberin is a hydrophobic polymer deposited as a secondary cell wall, that can be found in many plant organs such as potato periderm, seed coat, cork, root periderm and endodermis. The past two decades have seen extensive efforts to elucidate its biosynthesis from intracellular production of monomers to extracellular polymerization and deposition in suberin lamellae (48-52). Suberin plasticity in response to abiotic stresses such as drought, salt, waterlogging or cadmium, while observed in roots in many species, (20, 46, 47, 53), only recently started to be characterized at the molecular level. This topic gained increasing interest in the past few years after observing that endodermal suberin is even more plastic than previously thought, and not only overproduced in toxic environments but also tightly modulated in response to mineral deficiencies (6, 14, 21-25), to Casparian strip defects (9, 10, 12, 14, 16, 17) and during biotic interactions (25-28). In light of the plethora of signals controlling suberization, understanding the interaction between these pathways is critical. The potential interaction between ABA and SGN3/CIFs signaling has been previously interrogated (14, 36), suggesting complex coordination between root development and ABA-mediated responses as well as between roots and shoots to control suberization. Here, we demonstrate by pharmaco-genetic approaches that both pathways induce endodermal suberization independently (Fig. 1 and Fig. S1). Corroborating our conclusion, a recent large-scale approach (combining microbiome, ionome, and suberin analysis, and genetics) revealed that the plant microbiome influences suberization through suppression of ABA-mediated signaling but independent of the SGN3/CIFs pathway (25).

In our attempt to identify transcription factors that are involved in ABA- and/or SGN3/CIFs-mediated suberization, we expected to identify specific factors downstream of at least one of these two pathways. We benefited from the impressive work performed by the community in identifying MYB transcription factors sufficient to induce suberization (29-33) and found 4 MYB transcription factors (*MYB41, MYB53, MYB92* and *MYB93*) to be expressed in the endodermis at different degrees under unstressed conditions (Fig. 2B, Fig. 2C and Fig. 3C-E). To our surprise all of them are induced in the endodermis in response to both ABA and CIF2 application with *MYB41* and *MYB93* being expressed close to the root tip after both treatment (Fig. 2C, Fig. S2B-F, Fig. 3C-E, Fig. S3D-F). This suggests that these 4 MYBs form a point of convergence between ABA and SGN3/CIFs signaling in the endodermis, with the signal specificity being established upstream of *MYB41, MYB53, MYB92* and *MYB93*.

Confirming previous work in heterologous systems or whole-plant overexpression we found that these four MYBs are sufficient to induce ectopic suberization when strongly expressed one by one in the endodermis prior to suberization (state I of endodermal differentiation) (Fig. 2D-G, Fig S2G and Fig. 3B). Previous works showed *in vitro* that MYB41 can directly bind to the *LTP20* promoter (*LIPID TRANSFER PROTEIN20*; associated with functions in cutin and suberin export) (54) and MYB92 to the *BCCP2* promoter (*BIOTIN CARBOXYL CARRIER PROTEIN2*, involved in fatty acid synthesis) (33). Moreover, MYB53, MYB92 and MYB93 (as well as MYB9, MYB39 and MYB107) were shown in yeast one-hybrid and heterologous expression in tobacco leaves to activate the expression of *BCCP2* (33). In addition, MYB92 was shown to activate the expression of two other genes involved in fatty acid biosynthesis, *ACP1* (*ACYL CARRIER PROTEIN1*) and *LPD1* (*LIPOAMIDE DEHYDROGENASE1*) (33). Our analysis of conditional endodermal expression of *MYB41* showed that the expression of genes involved in suberin biosynthesis and polymerization is induced in roots shortly after MYB41 production (Fig 2G). Additionally, we showed that endodermal accumulation of MYB41 protein can trigger the expression of the suberin biosynthesis gene *GPAT5* shortly after (Fig. 2F, Fig. S2H-I). Unfortunately, despite multiple attempts we were unable to immunodetect MYB41 protein from roots (either using the MYB41-Venus version described in this study or by attempts to raise an anti-MYB41 antibody), that would have allowed us to identify its direct targets *in planta*. This is probably due to working in its endogenous tissue (the late differentiated endodermis) which represent comparatively few cells of a whole root combined with a low abundance of MYB41, the protein accumulating only transiently in few endodermal cells (Fig. 2F and Fig. S2I) . However, considering the high number of evidence from *in vitro*, yeast one-hybrid or transactivation assays in tobacco, we can hypothesize that most suberin-inducing MYBs, including the four MYBs of interest in this study (MYB41, MYB53, MYB92 and MYB93), could directly activate the expression not only of genes involved in the primary fatty acid biosynthesis but also suberin biosynthesis genes *in planta*.

Loss of function of single suberin-inducing *MYBs* was rarely undertaken. Phenotypes were described only for *myb9* and *myb107* mutants, whose seed coats display a reduction in suberin monomers and an increased permeability, and for *myb39* mutant displaying a reduction of suberin monomers in whole roots but only a minor delay of a few cells in endodermal suberization (31, 40, 41). The mutants *myb41, myb53* and *myb93* presented in this study, are not affected for suberin deposition in unstressed condition or in the presence of ABA or CIF2 (Fig. 3A,F). Interestingly the single mutant *myb92* displayed a significant delay in suberin deposition in unstressed condition but its suberin was still strongly induced in response to ABA and CIF2 to level similar to WT plants (Fig. 3F). We therefore took advantage of CRISPR/Cas9 gene editing to generate a quadruple *myb41-53-92-93* mutant (*quad-myb*). In non-stressed condition this *quad-myb* displayed a dramatic reduction of endodermal suberin with no suberin staining observed in endodermal cells and a reduction by 78% of suberin monomers detected in its roots (Fig. 4A, B, Fig. S4C). Mutants with such low amounts of endodermal suberin are extremely rare and most suberin biosynthesis mutants only moderately affect suberin amounts or its monomeric composition. For example, the *gpat5* mutant lacking a key enzyme for suberin biosynthesis displays only a 30% reduction of suberin monomer accumulating in its roots (55). To our knowledge, the only genotypes displaying a range of reduction comparable to the *quad-myb* is the quintuple *gelp22-38-49-51-96* mutant (affected in suberin polymerization in the cell wall), and the *ELTP::MYB4* line (where inhibition of the phenylpropanoid pathway in the endodermis leads to suberin detachment), both displaying an 85% reduction of suberin monomers in roots (43, 56). We are therefore confident that MYB41, MYB53, MYB92 and MYB93 form the core regulating machinery controlling suberization in the endodermis. However the slight differences in their expression territories with only *MYB41, MYB53* and *MYB92* expressed all along the suberizing zone and the suberin reduction observed in *myb92* but not in other single mutants suggest a certain level of specificity among these four *MYBs* in unstressed conditions (Fig. 2C, Fig. S2B, Fig. 3C-F). Importantly, in response to ABA or CIF2 the expression of all these *MYBs* were induced at different degrees resulting in all of them being highly expressed close to the root tip and all along the endodermis (Fig. 2C, Fig. S2B-C, Fig. 3C-E, Fig. S3D-F). Moreover, testing the effect of ABA, salt stress (previously shown to be ABA-dependent (6)), and CIF2, we found that suberin is virtually non-affected by these three treatments in *quad-myb* (Fig. 4A,E and Fig. S4C). MYB41, MYB53, MYB92 and MYB93 are therefore playing a central role in suberin induction by at least two independent signaling pathways. Yet, the fact that we could still observe in *quad-myb* a weak response (with few patches of suberized endodermal cells after ABA, salt or CIF2 treatment) (Fig. 4A and Fig. S4C), suggest that even more factors are either needed to fully regulate suberization or are not involved in suberization *per se* but capable to weakly compensate for the *quad-myb* defects. Such factors could be other endodermal *MYBs* with an endodermal expression induced by ABA and/or CIF2. Additionally, we could envision that other transcription factors such as bHLH transcription factor (basic Helix-Loop-Helix) and/or a WD40-repeat protein (WD, tryptophan-aspartic acid) could influence endodermal suberization. It is known that MYB-bHLH-WD40 protein complexes play central roles in controlling multiple cell fates such as root hair and trichome formation, anthocyanin biosynthesis, seed coat mucilage or pigmentation (57-59).

As outlined in the introduction, suberin function for plant nutrition has recently benefited from the identification of mutants and lines affected in endodermal suberization and the wide application of ionomic analysis (6, 8-11, 13, 25, 31). However, even though all these studies are of fundamental interest to unravel suberin function and its physiological relevance they are presently limited by the mutants and lines available at that time. In fact, plants with enhanced endodermal suberin, often characterized for their ionomic and physiological defects, are not specifically affected for this barrier. This is particularly the case for the enhanced suberin phenotypes being the consequence of Casparian strip defects, which activate SGN3/CIFs signaling and in turn lead to ectopic lignin and suberin deposition in the endodermis (9-12, 14, 16). In other words, the nutritional effect described in currently available analyses likely represent the consequence of multilevel defects in the endodermal barriers and of the activation of SGN3/CIF signaling. Because of these tissue-specific pleiotropic defects, the specific role played by suberin has remained unclear. On the other hand, mutants with a strong reduction of endodermal suberization were previously not available and studying a lack of suberin had been based on a synthetic line, artificially expressing a cutinase in the endodermis to degrade suberin (5, 6, 11, 14). While these lines showed a dramatic suberin reduction and were extremely important to distinguish between Casparian strip and suberin defects, we cannot exclude that artificially expressing the cutinase *CDEF1* in the endodermis would not lead to additional defects. Moreover, being highly plastic in response to nutrient availability (6, 14, 21-25), suberin defects described in non-stressed conditions can in some case be exacerbated or absent in stressed conditions (25). To fully understand suberin function in the endodermis we crucially need better and more specific mutants and lines with constitutively enhanced and reduced endodermal suberization. The lines presented here (*ELTP::MYB41* with constitutively enhanced suberization without any Casparian strip defects, and the *quad-myb* mutant with strongly reduced endodermal suberization and largely lacking regulation by ABA and salt stress) (Fig. 4A,B,E and Fig. S4C-E) provide such highly specific phenotypes. Their usefulness is highlighted by our ionomic analyses, which show clear differences between the enhanced suberin line *ELTP::MYB41*, a line combining enhanced suberin with Casparian strip defects (*CASP1::MYB41* line) (Fig. 4 D) and between the *quad-myb* and *ELTP::CDEF1* line (Fig. 4D). Importantly, the root development or expression of key genes involved in the acquisition of these elements being comparable between these lines these parameters are unlikely explaining the ionomic phenotypes observed (Fig. S5). In summary our results suggest that, in accordance with previous reports, suberin plays crucial roles for nutrient homeostasis, likely affecting directly transport through the endodermis. But its role might be more specific than initially thought, affecting mainly the acquisition of boron, sodium, calcium, arsenic and strontium in our experiments (Fig. 4D). We are therefore convinced that the tools generated in this study, especially the *quad-myb* and *ELTP::MYB41* plants will be of tremendous interest for the community in order to better understand suberin function in relation to nutrient availability as well as for its role in root development and biotic interactions. Given the increasing interest beyond fundamental research in manipulating suberin, extending the genetic tool box to specifically manipulate and fine-tune suberization is highly relevant for applied plant biology in crop improvement or carbon capture to combat climate change.

## Material and methods

### Plant material

All experiments were performed in Columbia-0 (Col-0) background. Previously published mutants and transgenic plants used in this study: *casp1-1 casp3-1*; *CASP1::NLS-GFP* (60); *esb1-1* (8); *sgn3-3, sgn3-4* (12); *ELTP::CDEF1*; *ELTP::NLS-3xmVenus* (6). The mutants *myb53* (SALK_076713), *myb92* (SM_3.41690) and *myb93* (SALK_131752) were obtained from NASC. Primers used for genotyping are presented Table S1. Transgenic lines previously described and slightly modified in this study *ELTP::abi1-1* (based on (4), here with a FastRed selection) *GPAT5::NLS-RFP* (based on *GPAT5::NLS-GFP* from (5)) and *GPAT5::mCitrine-SYP122* (6). The following mutants were generated for this study using *CRISPR*-Cas9 technology: *myb41_c1, myb41_c2* and *myb41_c2-myb53-myb92-myb93* (*quad-myb*, see constructs part for more details). The following transgenic lines were generated for this study: *MYB41::NLS-3xmVenus, MYB53::NLS-3xmVenus, MYB92::NLS-3xmVenus, MYB93::NLS-3xmVenus, GPAT5::NLS-3xmScarlet, CASP1xve::MYB41-mVenus, CASP1::MYB41, CASP1::MYB53, CASP1::MYB92, CASP1::MYB93, CASP1::myb41_c2, ELTP::MYB41, CASP1::myb53_c1, CASP1::myb92_c1* and *CASP1::myb93_c1*. The corresponding gene numbers are as follow: *CASP1*, At2g36100; *CASP3*, At2g27370; *ESB1*, At2g28670; *SGN3*, At4g20140; *CDEF1*, At4g30140; *ELTP*, At2g48140; *ABI1*, At4g26080; *GPAT5*, At3g11430; *SYP122*, At3g52400, *MYB41*, At4g28110; *MYB53*, At5g65230; *MYB92*, At5g10280; *MYB93*, At1g34670.

### Constructs

Plasmids were constructed using Multisite Gateway cloning (Thermo Fisher Scientific). The list of primers used for cloning are presented in Table S1. *MYB* promoter sequences upstream of ATG - *MYB41* (2167bp), *MYB53* (4117bp), *MYB92* (4098bp), *MYB93* (2873bp) were amplified from Col-0 genomic DNA and cloned into *pDONRP4-P1R* (Thermo Fisher Scientific). For promoter-reporter expression clones, *PROMOTER::NLS-3xmVenus or PROMOTER::NLS-3xmScarlet*, the entry plasmids containing the promoter region, along with *pDONRL1-NLS-3xmVenus-L2* (61) or *pDONRL1-NLS-3xmScarlet-L2* and the pEN-R2-*tNOS-* L3 containing the terminator *tNOS* (62) were recombined into the destination vectors pFR7m34GW or pFG7m34GW. The destination vectors pFR7m34GW or pFG7m34GW were obtained by substitution of the Hygromycin sequence in pH7m34GW by the FastRed and FastGreen sequences respectively (63). For endodermal specific expression of MYBs using *CASP1* and *ELTP* promoters (60, 64), MYB coding sequences were amplified from wild-type

Arabidopsis cDNA and cloned into pDONR221_L1-ORF-L2 vector were recombined with pDONR-P3-*tNOS-*P2R in the destination plasmid pFR7m34GW. Except for *MYB41cDNA* that was obtained from (29) and recloned into pDONR221_L1-CDDB-CAM-L2. For endodermal specific estradiol inducible *MYB41* expression (*CASP1xve::MYB41-mVenus)*, the entry vectors containing the inducible *CASP1* promoter pEN-L4-CASP1xve-R1 (65) was recombined with pDONR221_L1-MYB41nostop-L2 and pEN-R2-mVenus +stop-L3 into the destination vector pFG7m34GW. Cloning of vectors for CRISPR/Cas9 was done as previously described in (43, 44, 66). sgRNA for spCas9 were designed using webtools – CRISPR-P 2.0 design tool (http://crispr.hzau.edu.cn/CRISPR2/) (67) and Benchling (https://www.benchling.com). Pairs of annealed oligos of the sgRNA were cloned into the Bbs-I linearized entry vector (66) and recombined into the destination vector containing Cas9 expression cassette and a FastRed or FastGreen selection marker cassette. For large deletion of genomic regions in *myb41_c1* or multiplex targeting of *MYB53, MYB92* and *MYB93*, multiple entry vectors were used to clone different sgRNAs. Thereafter, recombined into the destination vector containing Cas9 expression cassette and FastRed or FastGreen selection marker cassette. After fluorescent seed selection in T1, non-fluorescent seeds in the T2 generation (indicating a segregation of the vector backbone containing the Cas9 cassette) were used to identify the mutations. Primary screening of mutants was done using High-Resolution Melting (HRM) curve analysis as previously described in (43). Candidates from HRM analysis were further confirmed for the mutations by sequencing of PCR-amplified genomic regions. Absence of off-target effects were controlled by sequencing the closest *MYB* homologues in the final mutant. To test the loss of function for *myb41_c2, myb53_c1, myb92*_c1 and *myb93*_*c1*, the corresponding cDNA were cloned from the mutated plants into pDONR221 and recombined with pEN-L4-CASP1-R1 and pDONR-P3-*tNOS-*P2R in the destination plasmid pFRm34GW. All constructs were transformed into Agrobacterium strain GV3101 by electroporation and used for transformation of Arabidopsis plants by the floral-dip method (68).

### Growth conditions

Seedlings for staining and live-imagine were grown vertically on square plates containing half-strength MS with 0.8% agar (Duchefa), without sucrose. Seeds were surface sterilized before sowing on plates and were incubated 2 to 4 days at 4°C and put to grow in growth chambers under continuous light (∼100 µE) at 22 °C. All histological and live-microscopy analysis were performed on 5-day old seedlings. For other experiments the age of the plant is specified in the figure legends. In soil, for amplifications and experiments in pots, plants were grown in long-day conditions (16 h day, 8 h night) with light intensity of 150-180 µE with 60-70% humidity and at 20 ± 2 °C.

### Pharmacological treatments

ABA hormone was stored at -20 °C in 50 mM stock solution, dissolved in methanol. For 16 h treatments, seedlings were transferred on solid half-MS media containing 1 µM ABA. For shorter treatments such as 3 h/6 h for staining, microscopy or gene expression analysis, stock solution of ABA was diluted to 1µM in liquid half-MS media applied directly on roots without transfer of seedlings. The peptide CIF2 described in (16, 19) (DY(sulfated)GHSSPKPKLVRPPFKLIPN) was stored as 1mM stock solution and the treatments with 1 µM were performed as described for the ABA treatment for short (3 h/6 h) and long (16 h) treatments. For 48 h fluridone treatments, 3-day seedlings were transferred on the half-MS containing 10 µM fluridone. Estradiol treatments were performed by diluting 5mM stock of estradiol to 5 µM in solid or liquid half-MS for long (16 h) short (3 h/6 h) respectively.

## Suberin staining

Whole mount suberin staining was performed as previously described in (6). Five-day old seedlings were incubated in Fluorol Yellow 088 (FY 088) (0.01% w/v, lactic acid) for 30 min at 70°C, washed twice with water and then counter-stained with Aniline Blue (0.5% w/v, water) for 30 min, washed with water and mounted on glass slide to be observed with an epifluorescence stereomicroscope-ZEISS Axio Zoom.V16 with a GFP filter ex: 450-490 nm, em: 500-550 nm. Samples were kept in the dark during the whole procedure. For subsequent suberin pattern quantifications, tiled images covering the whole seedlings in single images were captured. For imaging of the large field of view with high-resolution, multiple smaller images were captured as tiles and stitched. Region of interest of the root was defined by marking the ‘tile-region’ after a quick scan of the sample at lower resolution. Adequate number of focus points were used to adjust the focus of the sample along the region of interest. 10% area of overlap was defined for alignment and stitching of tiles. Fiji (http://fiji.sc/Fiji) (69) was used on Zen2.3 blue exported stitched tile images for quantification of suberin patterns (in mm) along the root: suberized for the fully suberized zone, patchy for the partially suberized zone and non-suberized –for the zone prior to suberization. Results are presented as percentage of the root as previously done (6, 12).

### Lignin staining

CearSee-adapted cell wall staining was performed as described (70). Briefly, 5-day-old seedlings were fixed in 1 × PBS containing 4% paraformaldehyde, 1 h at room temperature and washed twice with 1 × PBS. Following fixation, the seedlings were cleared overnight in ClearSee solution after which the solution was exchanged to 0.2% Basic Fuchsin in ClearSee solution lignin staining. After overnight staining, the dye solution was removed and rinsed once with ClearSee solution, the seedlings were subsequently washed in ClearSee solution for 30 min and washed again in another ClearSee solution for at least one overnight before observation with a Leica SP8 confocal. All clearing, staining and washing steps were performed in 12 well plates, covered with aluminum foil and under gentle agitation.

### Propidium iodide test

Propidium iodide (PI) was used as an apoplastic tracer to assess Casparian strip functionality as previously described (5, 65). Seedlings were live-stained with 15 µM PI; kept in the dark for 10 min and then rinsed twice with water. The apoplastic barrier was determined under a fluorescent Leica DM6 B microscope with I3 filter and 20x magnification, as the number of endodermal cells after the onset of elongation where PI uptake is blocked at the endodermis. The onset of elongation was defined as the first endodermal cell for which the length was at least three times its width.

### Confocal microscopy

Confocal laser scanning microscopy experiments were performed either on a Zeiss LSM 780, a Zeiss LSM 800 or a Leica SP8 microscopes. Excitation and detection windows were set as follows: Zeiss LSM 780: mVenus ex: 488 nm, em: 519-559 nm; RFP/mScarlet ex: 543 nm, em: 591-637 nm ; Zeiss LSM 800: mCITRINE/mVenus ex: 488 nm, em: 500-546 nm; RFP/mScarlet ex: 561 nm, em: 585-617 nm; PI ex: 561 nm, em: 592-617 nm ; Leica SP8: Basic Fuchsin ex: 561 nm, em: 600-650 nm. For imaging of the large field of view with high-resolution, multiple smaller images were captured as tiles and stitched together for a larger view of roots. Region of interest of the root was defined by marking the ‘tile-region’ after a quick scan of the sample at lower resolution. Adequate number of focus points were used to adjust the focus of the sample along the region of interest. Acquisition of tiled images was combined with Z-stacking and in certain cases with time series as well. 10% area of overlap was defined for alignment and stitching of tiles and tiled Z-stacks were used for orthogonal projection and subsequently exported. For time-course experiments, 25-30 min time interval in between the scans was defined for 10-12 cycles. Scanner and detector settings were kept unchanged for every experiment. Images were analyzed with Zen2.3 blue (LSM 800) or Zen2.3 black (LSM 780) software and Fiji (http://fiji.sc/Fiji) (69). Fluorescence intensities were calculated nucleus by nucleus along one cell file from the onset of nuclear signal, considering the maximum intensity detected in each individual nucleus as an estimate the difference of intensity between nuclei.

### Q-RT-PCR

For gene expression analysis, 25-30 roots of 7d-ay-old seedlings were harvested and pooled together to form one biological replicate. RNA extractions were performed by Trizol-adapted RNeasy MinElute Cleanup Kit (Qiagen). RNA was reverse-transcribed using Thermo Scientific Maxima First Strand cDNA Synthesis Kit following the manufacturer’s protocol. Real-time PCR was performed on Applied Biosystems QuantStudio5 thermocycler using

Applied Biosystems SYBR Green master mix. *ACTIN-2 (At3g18780)* was used as the housekeeping gene and relative expression of each gene was calculated using the 2^-^ΔΔCt method (71). The list of primer used for Q-RT-PCR are presented in Table S2.

### Chemical suberin analysis

We used the protocol as described by (72) for the analysis of ester-bound lipids, which likely only belong suberin in the described organ and developmental stage. In brief, 200 mg of seeds were grown and after five days, the roots (between 200 and 300 per replicate) were shaved off after flash freezing and extracted in isopropanol/0.01% butylated hydroxytoluene (BHT). They were then delipidized two times (16h, 8h) in each of the following solvents, i.e., chloroform-methanol (2:1), chloroform-methanol (1:1), methanol each with 0.01% BHT, under agitation before being dried for 3 days under vacuum. Depolymerization was performed by base catalysis (73). Briefly, dried plant samples were trans-esterified in 2 mL of reaction medium. 20 mL reaction medium was composed of 3 mL methyl acetate, 5 mL of 25% sodium methoxide in dry methanol and 12 mL dry methanol. The equivalents of 5 mg of methyl heptadecanoate and 10 mg of ω-pentadeca-lactone/sample were added as internal standards. After incubation of the samples at 60°C for 2h 3.5 mL dichloromethane, 0.7 mL glacial acetic acid and 1 mL 0.9% NaCl (w/v) /100 mM Tris-HCl (pH 8.0) were added to each sample and subsequently vortexed for 20 s. After centrifugation (1500g for 2 min), the organic phase was collected, washed with 2 mL of 0.9% NaCl, and dried over sodium sulfate.

The organic phase was then recovered and concentrated under a stream of nitrogen. The resulting suberin monomer fraction was derivatized with BFTSA/pyridine (1:1) at 70°C for 1 h and injected out of hexane on a HP-5MS column (J&W Scientific) in a gas chromatograph coupled to a mass spectrometer and a flame ionization detector (Agilent 6890N GC Network systems). The temperature cycle of the oven was the following: 2 min at 50°C, increment of 20°C/min to 160°C, of 2°C/min to 250°C and 10°C/min to 310°C, held for 15 min. 3 independent experiments were performed with 4 replicates for each genotype, respectively, and a representative dataset is presented. The amounts of unsubstituted C16 and C18 fatty acids were not evaluated because of their omnipresence in the plant and in the environment.

### Ionomic analysis

Leaf elemental content was measured using ICP-MS as previously described (74). Briefly, dried leaves were transferred into the Pyrex test tubes, weighted, and digested with 1 ml of concentrated trace metal grade nitric acid Primar Plus (Fisher Chemicals) containing an indium internal standard, in the dry block heaters (SCP Science; QMX Laboratories) at 115°C for 4 h. After cooling, digested samples were diluted to 10mL with 18.2 MΩcm Milli-Q Direct water (Merck Millipore) and elemental analysis was performed using an ICP-MS (PerkinElmer NexION 2000 equipped with Elemental Scientific Inc autosampler) in the collision mode (He). Twenty-three elements were monitored (Li, B, Na, Mg, P, S, K, Ca, Mn, Fe, Co, Ni, Cu, Zn, As, Rb, Sr, Mo and Cd). A matrix-matched liquid reference material composed of pooled digested samples was prepared before the beginning of the sample run and used every ninth sample to correct for variation within ICP-MS analysis runs. The calibration standards were prepared from single element standards solutions (Inorganic Ventures; Essex Scientific Laboratory Supplies Ltd, Essex, UK). Samples concentrations were calculated using external calibration method within the instrument software. The final concentrations were obtained by normalizing the element concentrations to the sample dry weight.

### Statistical analysis

Statistical analyses were done with the GraphPad Prism 8.0 software (https://www.graphpad.com/) or with the R environment (75). For statistical analysis of multiple transgenic lines, genotypes or treatments parametric or nonparametric One-way or Two-way ANOVA, and Tukey’s test were used as a multiple comparison procedures. Binary comparisons were performed using Student’s t-test. Statistical representation for specific experiment are described in figure legends. The data are presented as mean ± standard deviation, and “n” represents number of biological replicates.

## Acknowledgments and funding sources

We would like to thank Robertas Ursache for sharing plasmids and methods for CRISPR/Cas9 mutagenesis and Lothar Kalmbach for destination plasmids allowing Fastgreen and Fastred selection after triple Gateway cloning and critical reading of the manuscript. Owen Rowland and Dylan Kosma are thanked for sharing the MYB41 cDNA and Michael Hothorn for sharing CIF2 peptide. We are thankful for Sylvain Loubéry and the Bioimaging center at the University of Geneva for assistance with fluorescent microscopy and to Rochus B. Franke for critical input on chemical analysis for suberin. We would also like to thank Niko Geldner and his lab where preliminary work for this project was conducted. This work was supported by funding from the Sandoz Family Monique De Meuron philanthropic foundation’s program for academic promotion and the SNSF (grant numbers 31003A_179159 and project number PCEGP3_187007 to MB and grant number 310030_188672 to CN).

## Notes

### Competing Interest Statement

The authors have declared no competing interest.

## References

1. Barberon M (2017) The endodermis as a checkpoint for nutrients. New Phytologist 213(4):1604–1610.

2. Ramakrishna P & Barberon M (2019) Polarized transport across root epithelia. Curr Opin Plant Biol 52:23–29.

3. Alassimone J, Naseer S, & Geldner N (2010) A developmental framework for endodermal differentiation and polarity. Proceedings of the National Academy of Sciences 107(11):5214–5219.

4. Barbosa ICR, Rojas-Murcia N, & Geldner N (2019) The Casparian strip-one ring to bring cell biology to lignification? Current opinion in biotechnology 56:121–129.

5. Naseer S, et al. (2012) Casparian strip diffusion barrier in Arabidopsis is made of a lignin polymer without suberin. Proceedings of the National Academy of Sciences 109(25):10101–10106.

6. Barberon M, et al. (2016) Adaptation of Root Function by Nutrient-Induced Plasticity of Endodermal Differentiation. Cell 164(3):447–459.

7. Robbins II NE, Trontin C, Duan L, & Dinneny JR (2014) Beyond the barrier: communication in the root through the endodermis. Plant Physiology 166(2):551– 559.

8. Baxter I, et al. (2009) Root Suberin Forms an Extracellular Barrier That Affects Water Relations and Mineral Nutrition in Arabidopsis. PLoS Genet 5(5):e1000492.

9. Hosmani PS, et al. (2013) Dirigent domain-containing protein is part of the machinery required for formation of the lignin-based Casparian strip in the root. Proceedings of the National Academy of Sciences 110(35):14498–14503.

10. Kamiya T, et al. (2015) The MYB36 transcription factor orchestrates Casparian strip formation. Proceedings of the National Academy of Sciences 112(33):10533–10538.

11. Li B, et al. (2017) Role of LOTR1 in Nutrient Transport through Organization of Spatial Distribution of Root Endodermal Barriers. Current Biology 27(5):758–765.

12. Pfister A, et al. (2014) A receptor-like kinase mutant with absent endodermal diffusion barrier displays selective nutrient homeostasis defects. Elife 3.

13. Wang Z, et al. (2019) OsCASP1 Is Required for Casparian Strip Formation at Endodermal Cells of Rice Roots for Selective Uptake of Mineral Elements. Plant Cell 31(11):2636–2648.

14. Wang P, et al. (2019) Surveillance of cell wall diffusion barrier integrity modulates water and solute transport in plants. Scientific reports 9(1):4227.

15. Doblas VG, Geldner N, & Barberon M (2017) The endodermis, a tightly controlled barrier for nutrients. Curr Opin Plant Biol 39:136–143.

16. Doblas VG, et al. (2017) Root diffusion barrier control by a vasculature-derived peptide binding to the SGN3 receptor. Science 355(6322):280–284.

17. Fujita S, et al. (2020) SCHENGEN receptor module drives localized ROS production and lignification in plant roots. The EMBO journal 39(9):e103894.

18. Nakayama T, et al. (2017) A peptide hormone required for Casparian strip diffusion barrier formation in Arabidopsis roots. Science 355(6322):284–286.

19. Okuda S, et al. (2020) Molecular mechanism for the recognition of sequence-divergent CIF peptides by the plant receptor kinases GSO1/SGN3 and GSO2. Proc Natl Acad Sci U S A 117(5):2693–2703.

20. Líška D, Martinka M, Kohanová J, & Lux A (2016) Asymmetrical development of root endodermis and exodermis in reaction to abiotic stresses. Annals of Botany 118(4):667–674.

21. Chen A, Husted S, Salt DE, Schjoerring JK, & Persson DP (2019) The Intensity of Manganese Deficiency Strongly Affects Root Endodermal Suberization and Ion Homeostasis. Plant Physiol 181(2):729–742.

22. Knipfer T, Danjou M, Vionne C, & Fricke W (2020) Salt stress reduces root water uptake in barley (Hordeum vulgare L.) through modification of the transcellular transport path. Plant Cell Environ 44(2):458–475.

23. Namyslov J, Bauriedlová Z, Janoušková J, Soukup A, & Tylová E (2020) Exodermis and Endodermis Respond to Nutrient Deficiency in Nutrient-Specific and Localized Manner. Plants (Basel, Switzerland) 9(2):201.

24. Armand T, Cullen M, Boiziot F, Li L, & Fricke W (2019) Cortex cell hydraulic conductivity, endodermal apoplastic barriers and root hydraulics change in barley (Hordeum vulgare L.) in response to a low supply of N and P. Ann Bot 124(6):1091– 1107.

25. Salas-González I, et al. (2021) Coordination between microbiota and root endodermis supports plant mineral nutrient homeostasis. Science 371(6525):eabd0695.

26. Holbein J, et al. (2019) Root endodermal barrier system contributes to defence against plant-parasitic cyst and root-knot nematodes. Plant J 100(2):221–236.

27. Emonet A, et al. (2021) Spatially Restricted Immune Responses Are Required for Maintaining Root Meristematic Activity upon Detection of Bacteria. Current Biology 31(5):1012–1028.e1017.

28. Fröschel C, et al. (2021) Plant roots employ cell-layer-specific programs to respond to pathogenic and beneficial microbes. Cell Host & Microbe 29(2):299–310.e297.

29. Kosma DK, et al. (2014) AtMYB41 activates ectopic suberin synthesis and assembly in multiple plant species and cell types. The Plant Journal 80(2):216–229.

30. Wei X, et al. (2019) Three Transcription Activators of ABA Signaling Positively Regulate Suberin Monomer Synthesis by Activating Cytochrome P450 CYP86A1 in Kiwifruit. Front Plant Sci 10:1650.

31. Cohen H, Fedyuk V, Wang C, Wu S, & Aharoni A (2020) SUBERMAN Regulates Developmental Suberization of the Arabidopsis Root Endodermis. Plant J 102(3):431– 447.

32. Mahmood K, et al. (2019) Overexpression of ANAC046 Promotes Suberin Biosynthesis in Roots of Arabidopsis thaliana. International journal of molecular sciences 20(24):6117.

33. To A, et al. (2020) AtMYB92 enhances fatty acid synthesis and suberin deposition in leaves of Nicotiana benthamiana. Plant J 103(2):660–676.

34. Gibbs DJ, et al. (2014) AtMYB93 is a novel negative regulator of lateral root development in Arabidopsis. The New phytologist 203(4):1194–1207.

35. Andersen TG, et al. (2018) Diffusible repression of cytokinin signalling produces endodermal symmetry and passage cells. Nature 555(7697):529–533.

36. Wang C, et al. (2020) Developmental programs interact with abscisic acid to coordinate root suberization in Arabidopsis. Plant J 104(1):241–251.

37. Popova LP & Riddle KA (1996) Development and accumulation of ABA in fluridone-treated and drought-stressed Vicia faba plants under different light conditions. Physiologia Plantarum 98(4):791–797.

38. Stewart CR & Voetberg G (1987) Abscisic Acid Accumulation Is Not Required for Proline Accumulation in Wilted Leaves. Plant Physiology 83(4):747–749.

39. Xu N & Bewley JD (1995) The role of abscisic acid in germination, storage protein synthesis and desiccation tolerance in alfalfa (Medicago sativa L.) seeds, as shown by inhibition of its synthesis by fluridone during development. Journal of Experimental Botany 46(6):687–694.

40. Lashbrooke J, et al. (2016) MYB107 and MYB9 Homologs Regulate Suberin Deposition in Angiosperms. Plant Cell 28(9):2097–2116.

41. Gou M, et al. (2017) The MYB107 Transcription Factor Positively Regulates Suberin Biosynthesis. Plant Physiology 173(2):1045–1058.

42. Goda H, et al. (2008) The AtGenExpress hormone and chemical treatment data set: experimental design, data evaluation, model data analysis and data access. The Plant Journal 55(3):526–542.

43. Ursache R, et al. (2021) GDSL-domain proteins have key roles in suberin polymerization and degradation. Nature Plants 7(3):353–364.

44. Ursache R, Fujita S, Tendon VD, & Geldner N (2021) Combined fluorescent seed selection and multiplex CRISPR/Cas9 assembly for fast generation of multiple Arabidopsis mutants. bioRxiv:2021.2005.2020.444986.

45. Krishnamurthy P, et al. (2020) Regulation of a Cytochrome P450 Gene CYP94B1 by WRKY33 Transcription Factor Controls Apoplastic Barrier Formation in Roots to Confer Salt Tolerance. Plant Physiol 184(4):2199–2215.

46. Krishnamurthy P, et al. (2009) The role of root apoplastic transport barriers in salt tolerance of rice (Oryza sativa L.). Planta 230(1):119–134.

47. Krishnamurthy P, Ranathunge K, Nayak S, Schreiber L, & Mathew MK (2011) Root apoplastic barriers block Na+ transport to shoots in rice (Oryza sativa L.). Journal of Experimental Botany 62(12):4215–4228.

48. Beisson F, Li-Beisson Y, & Pollard M (2012) Solving the puzzles of cutin and suberin polymer biosynthesis. Curr Opin Plant Biol 15(3):329–337.

49. Philippe G, et al. (2020) Cutin and suberin: assembly and origins of specialized lipidic cell wall scaffolds. Curr Opin Plant Biol 55:11–20.

50. Graça J (2015) Suberin: the biopolyester at the frontier of plants. Frontiers in chemistry 3:62.

51. Vishwanath SJ, Delude C, Domergue F, & Rowland O (2015) Suberin: biosynthesis, regulation, and polymer assembly of a protective extracellular barrier. Plant cell reports 34(4):573–586.

52. Ranathunge K, Schreiber L, & Franke R (2011) Suberin research in the genomics era— New interest for an old polymer. Plant Science 180(3):399–413.

53. Shiono K, et al. (2014) RCN1/OsABCG5, an ATP-binding cassette (ABC) transporter, is required for hypodermal suberization of roots in rice (Oryza sativa). Plant Journal 80(1):40–51.

54. Hoang MH, et al. (2012) Phosphorylation by AtMPK6 is required for the biological function of AtMYB41 in Arabidopsis. Biochemical and biophysical research communications 422(1):181–186.

55. Beisson F, Li YH, Bonaventure G, Pollard M, & Ohlrogge JB (2007) The acyltransferase GPAT5 is required for the synthesis of suberin in seed coat and root of Arabidopsis. Plant Cell 19(1):351–368.

56. Andersen TG, et al. (2020) Tissue-autonomous phenylpropanoid production is essential for establishment of root barriers. bioRxiv:2020.2006.2018.159475.

57. Robinson DO & Roeder AHK (2015) Themes and variations in cell type patterning in the plant epidermis. Current Opinion in Genetics & Development 32:55–65.

58. Xu W, Dubos C, & Lepiniec L (2015) Transcriptional control of flavonoid biosynthesis by MYB-bHLH-WDR complexes. Trends Plant Sci 20(3):176–185.

59. Millard PS, Weber K, Kragelund BB, & Burow M (2019) Specificity of MYB interactions relies on motifs in ordered and disordered contexts. Nucleic Acids Research 47(18):9592–9608.

60. Roppolo D, et al. (2011) A novel protein family mediates Casparian strip formation in the endodermis. Nature 473(7347):380–383.

61. Robe K, et al. (2021) Coumarin accumulation and trafficking in Arabidopsis thaliana: a complex and dynamic process. The New phytologist 229(4):2062–2079.

62. Karimi M, Bleys A, Vanderhaeghen R, & Hilson P (2007) Building blocks for plant gene assembly. Plant Physiol 145(4):1183–1191.

63. Shimada TL, Shimada T, & Hara-Nishimura I (2010) A rapid and non-destructive screenable marker, FAST, for identifying transformed seeds of Arabidopsis thaliana. Plant J 61(3):519–528.

64. Wyrsch I, Domínguez-Ferreras A, Geldner N, & Boller T (2015) Tissue-specific FLAGELLIN-SENSING 2 (FLS2) expression in roots restores immune responses in Arabidopsis fls2 mutants. New Phytologist 206(2):774–784.

65. Lee Y, Rubio MC, Alassimone J, & Geldner N (2013) A Mechanism for Localized Lignin Deposition in the Endodermis. Cell 153(2):402–412.

66. Fauser F, Schiml S, & Puchta H (2014) Both CRISPR/Cas-based nucleases and nickases can be used efficiently for genome engineering in Arabidopsis thaliana. The Plant Journal 79(2):348–359.

67. Liu H, et al. (2017) CRISPR-P 2.0: An Improved CRISPR-Cas9 Tool for Genome Editing in Plants. Molecular Plant 10(3):530–532.

68. Clough SJ & Bent AF (1998) Floral dip: a simplified method forAgrobacterium-mediated transformation ofArabidopsis thaliana. The Plant Journal 16(6):735–743.

69. Schindelin J, et al. (2012) Fiji: an open-source platform for biological-image analysis. Nature Methods 9(7):676–682.

70. Ursache R, Andersen TG, Marhavý P, & Geldner N (2018) A protocol for combining fluorescent proteins with histological stains for diverse cell wall components. Plant J 93(2):399–412.

71. Livak KJ & Schmittgen TD (2001) Analysis of relative gene expression data using real-time quantitative PCR and the 2(-Delta Delta C(T)) Method. Methods (San Diego, Calif.) 25(4):402–408.

72. Berhin A, et al. (2019) The Root Cap Cuticle: A Cell Wall Structure for Seedling Establishment and Lateral Root Formation. Cell 176(6):1367–1378.e1368.

73. Li-Beisson Y, et al. (2013) Acyl-Lipid Metabolism. The Arabidopsis Book 2013(11).

74. Danku JM, Lahner B, Yakubova E, & Salt DE (2013) Large-scale plant ionomics. Methods in molecular biology (Clifton, N.J.) 953:255–276.

75. Team RC (2019) R: A language and environment for statistical computing.

