## Supplementary figures and images for "Suberin plasticity to developmental and exogenous cues is regulated by a set of MYB transcription factors"

### Fig. S2

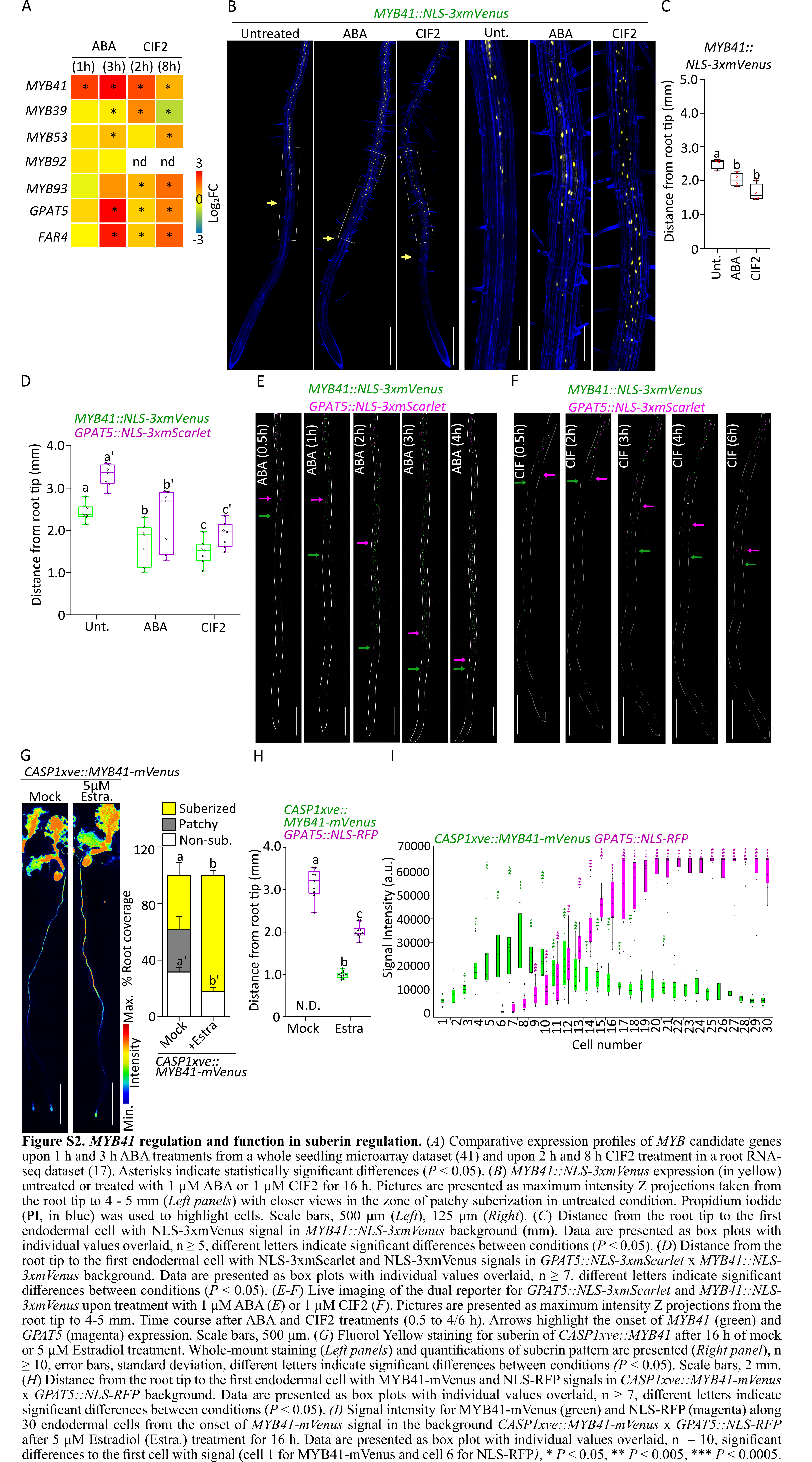

### Fig. S3

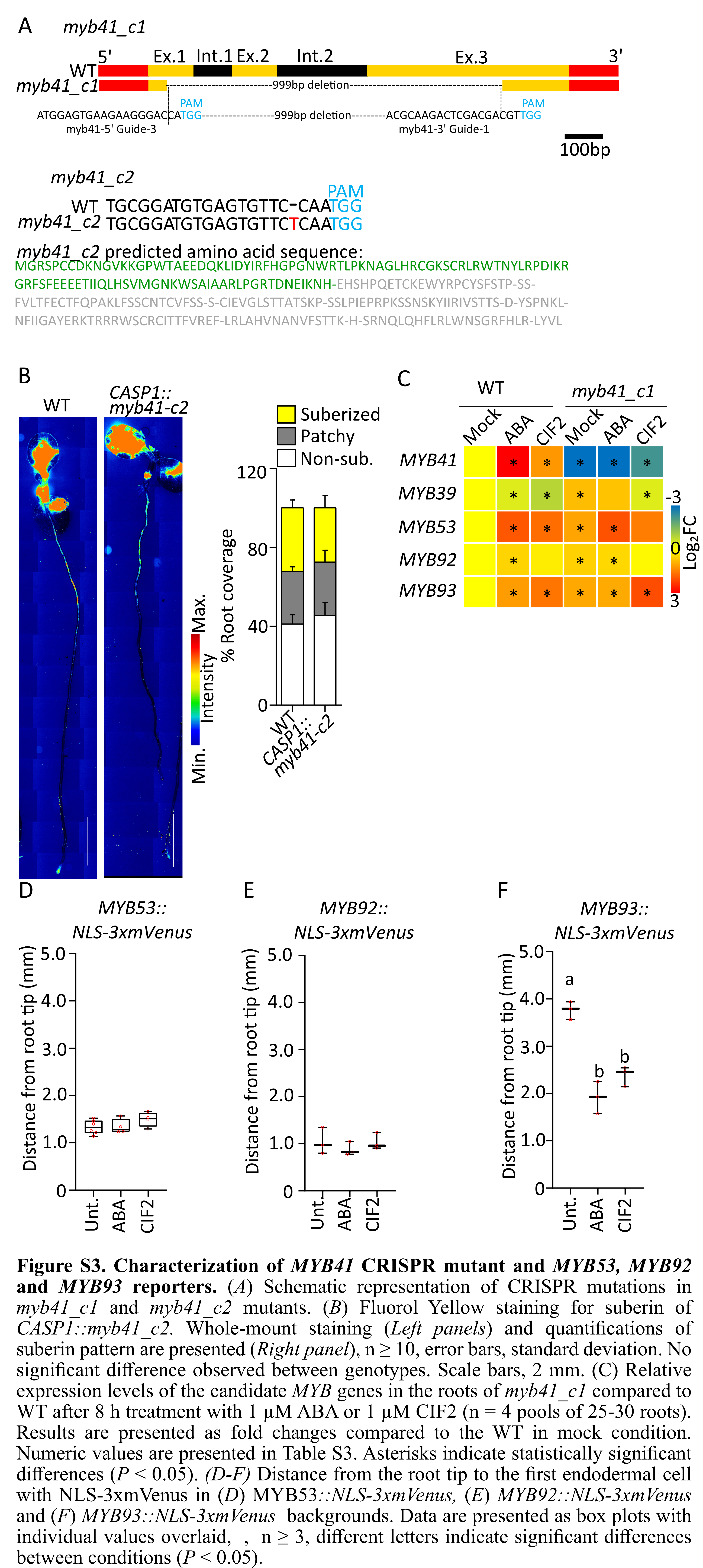

### Fig. S4

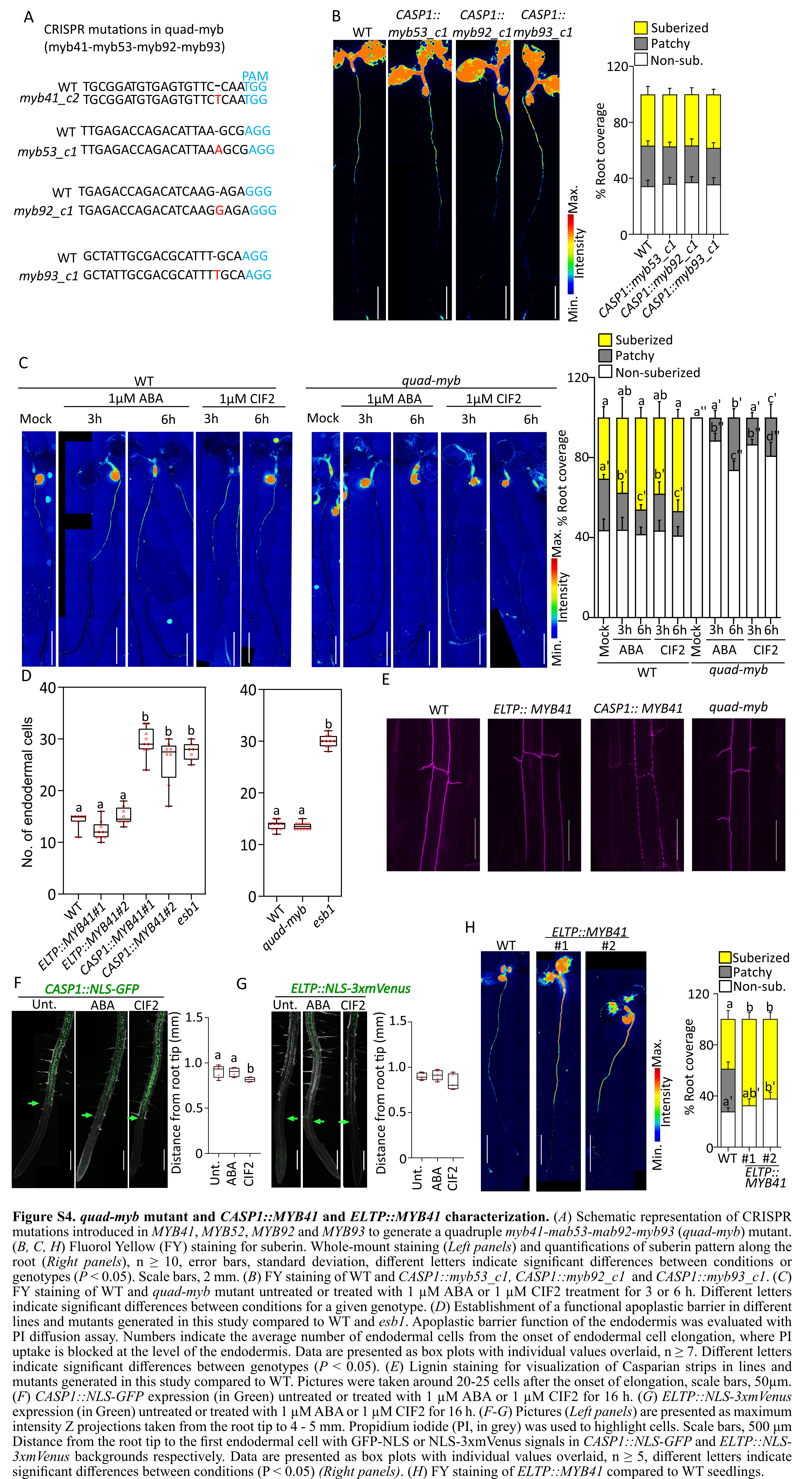

### Fig. S5

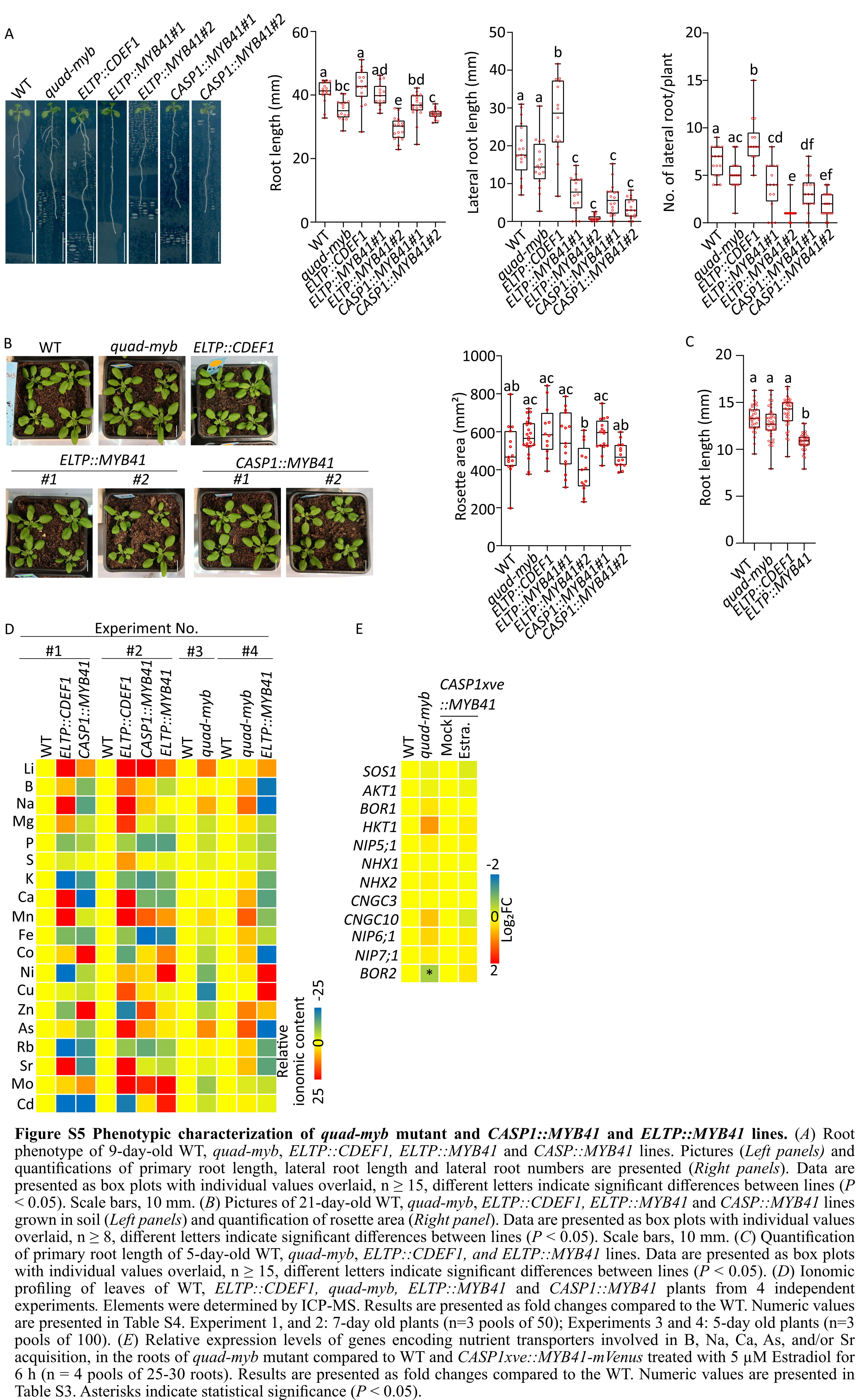

### Fig.S1

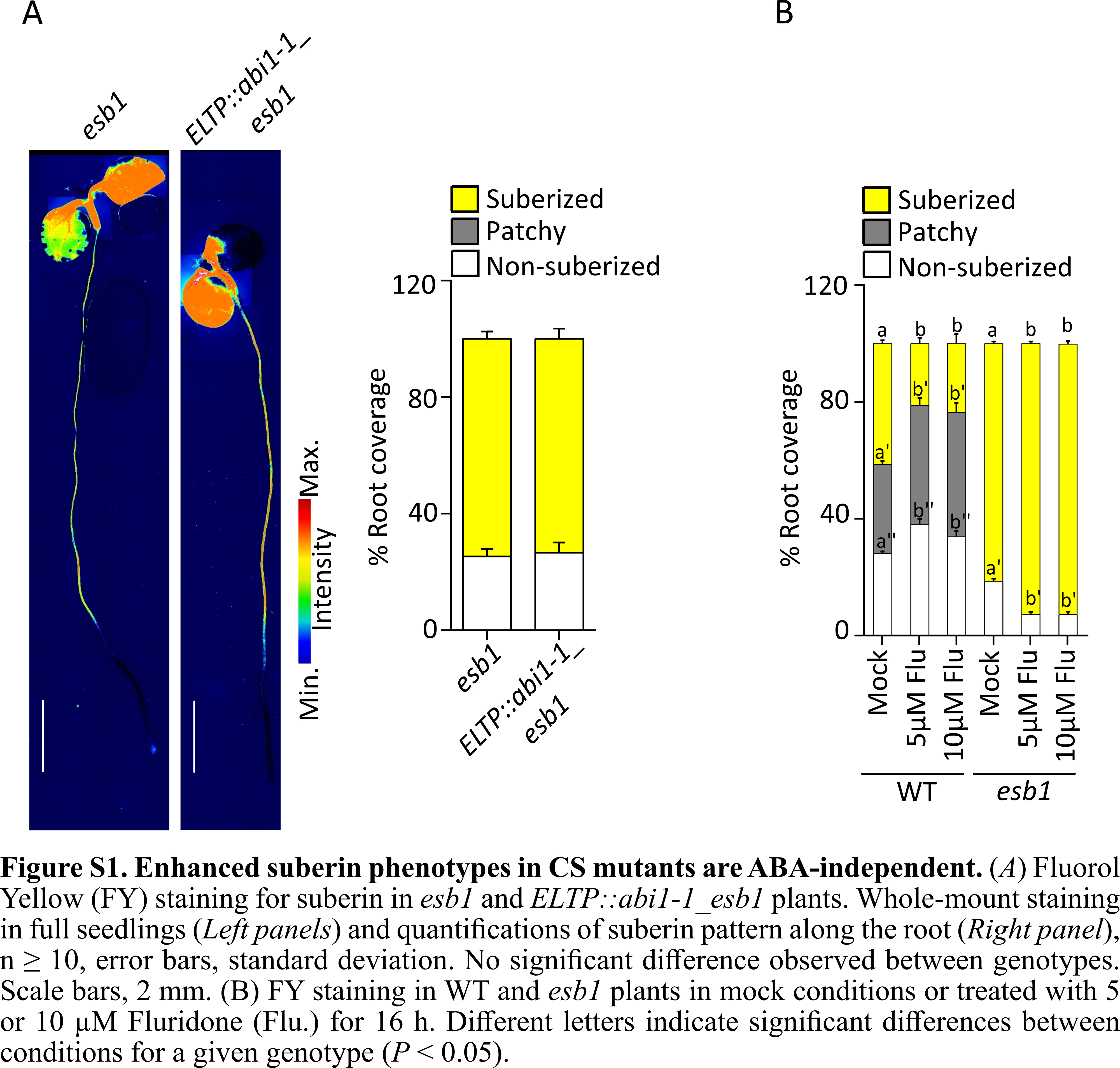

### Table S1

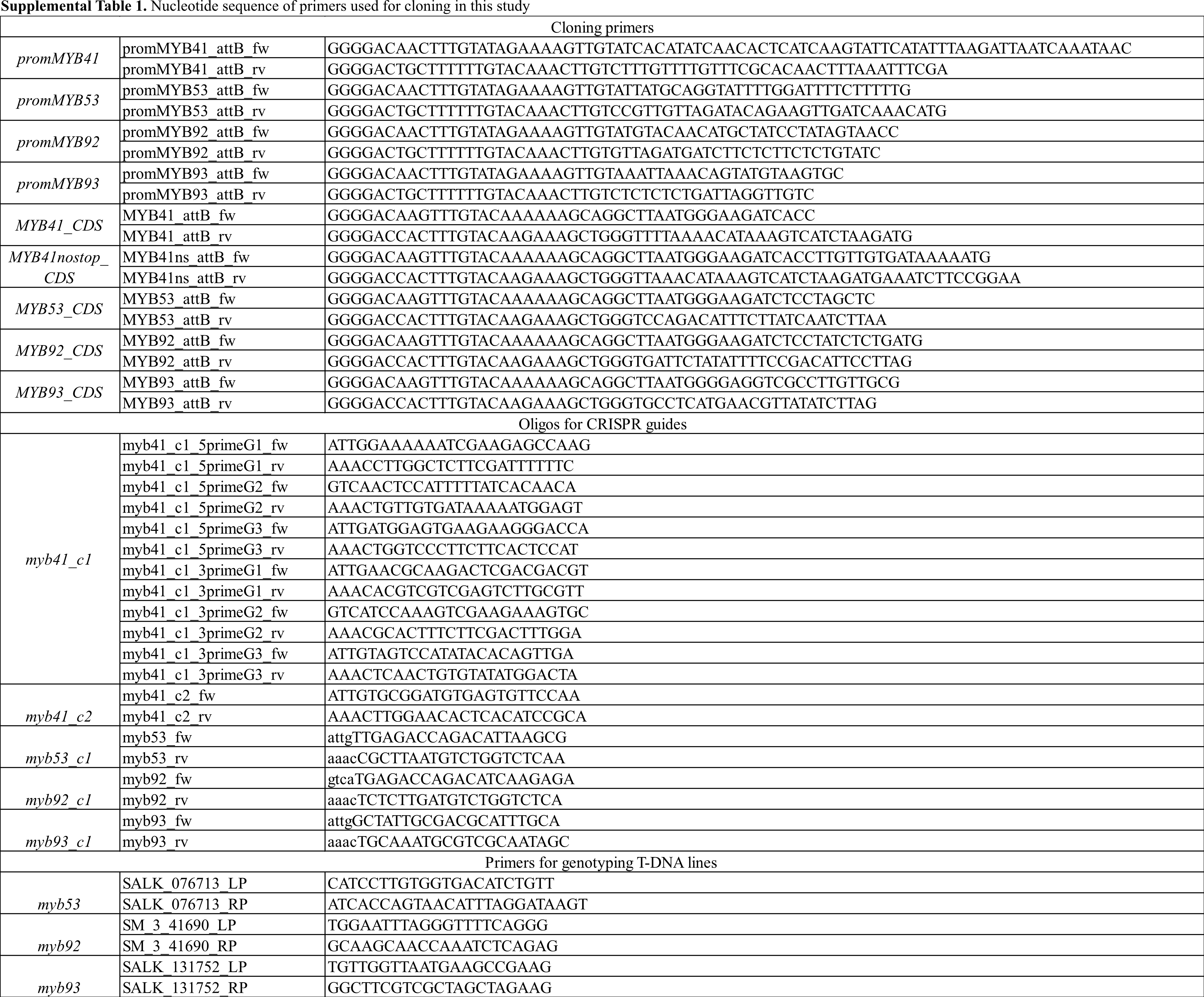

### Table S2

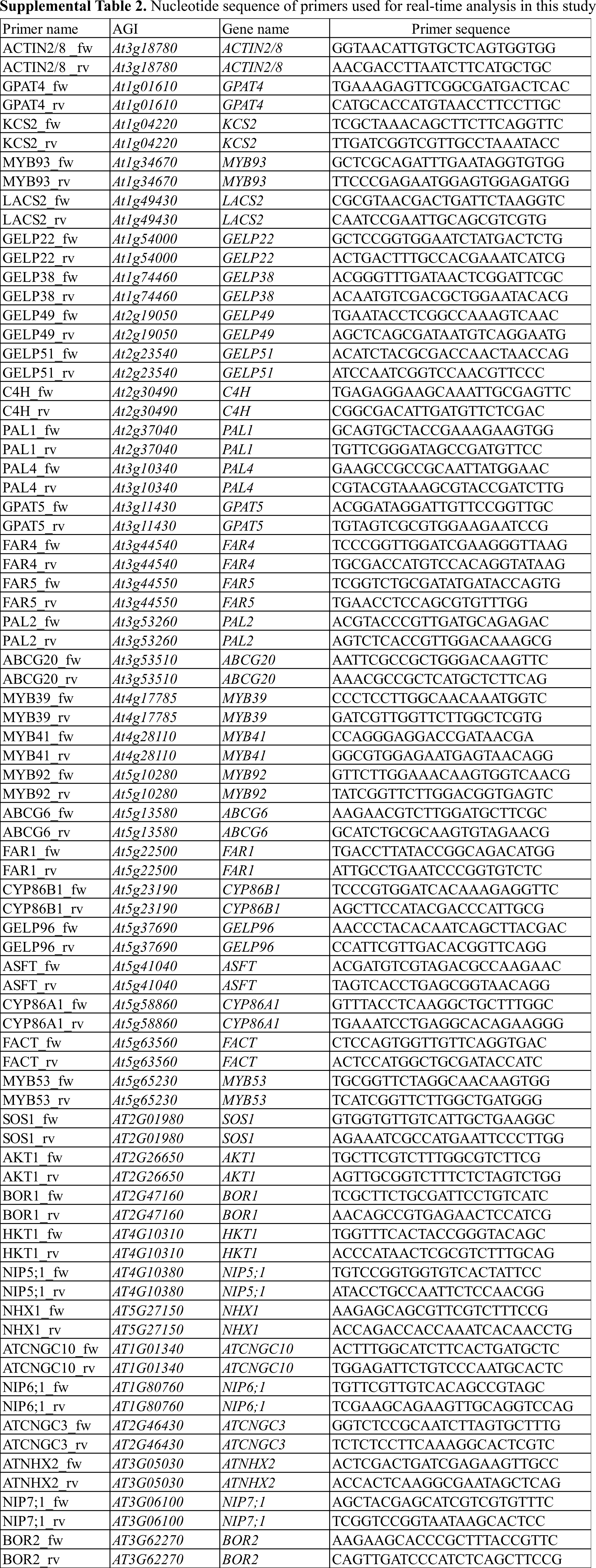

### Table S3A

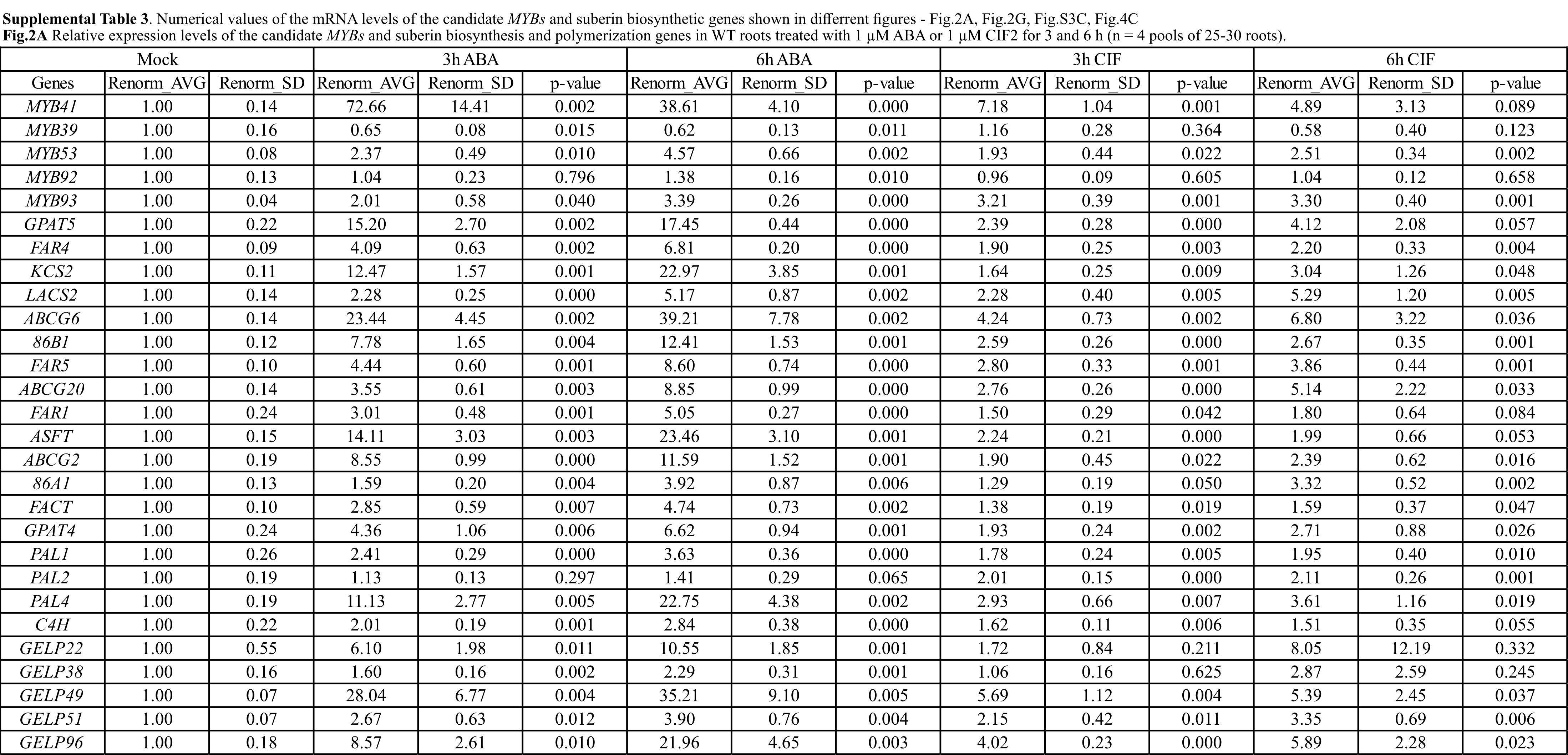

### Table S3B

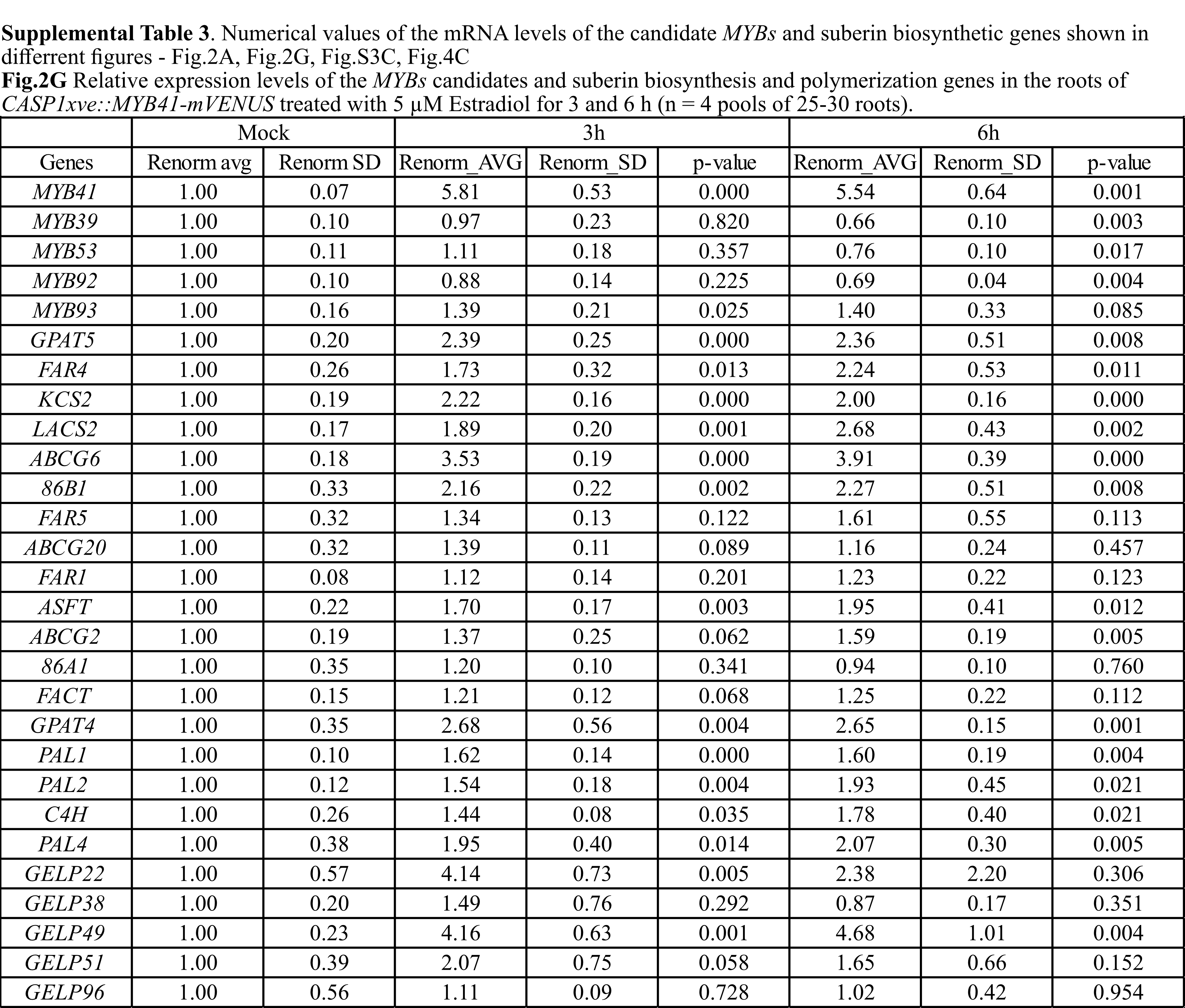

### Table S3C

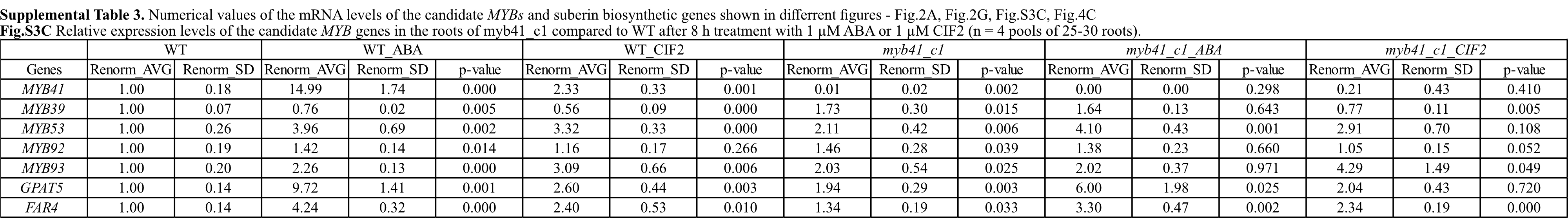

### Table S3D

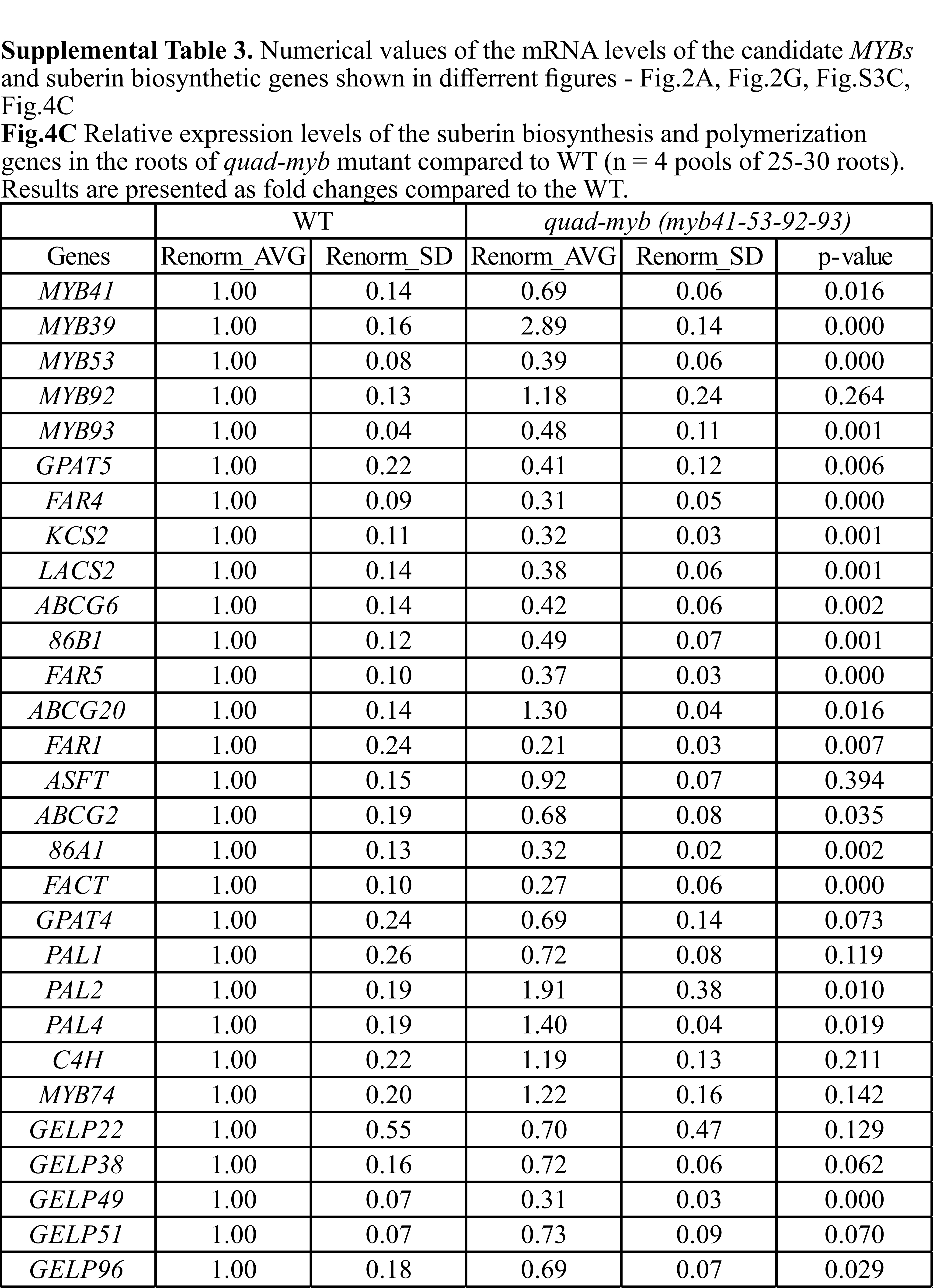

### Table S3E

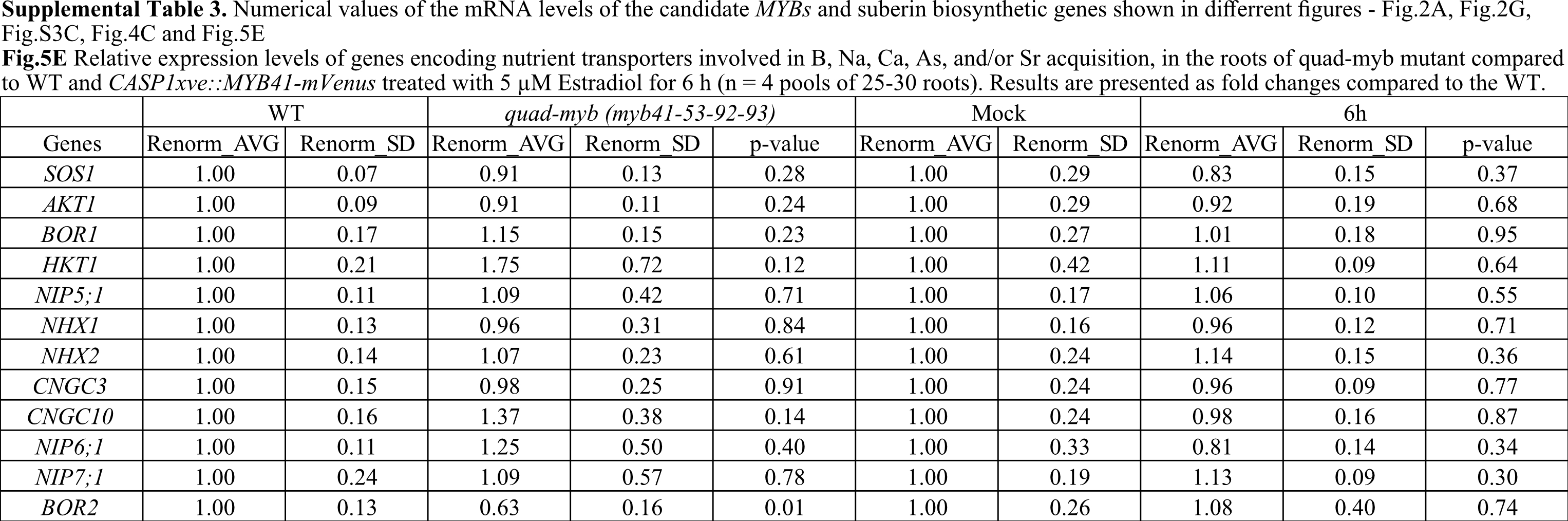

### Table S4

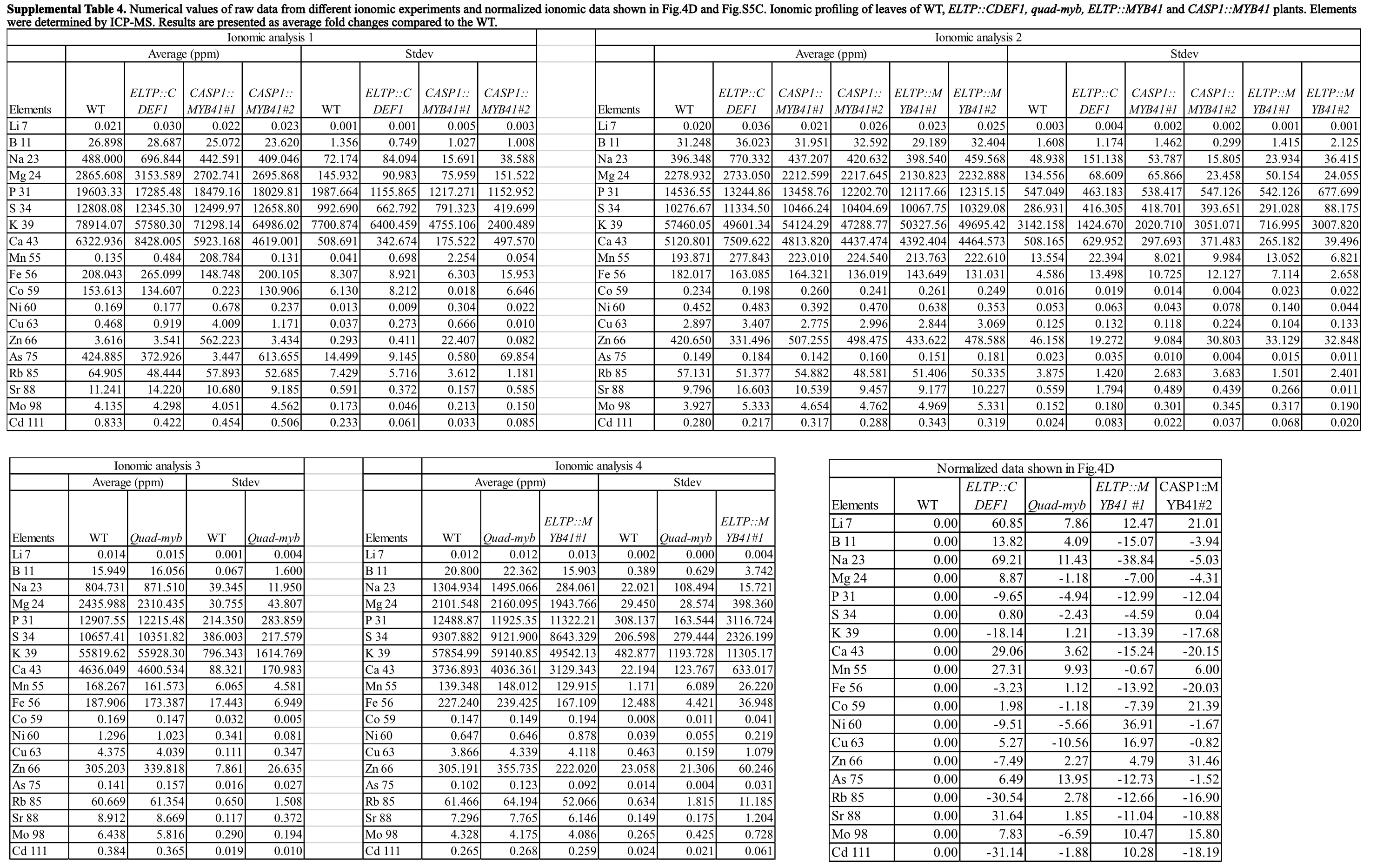
